# Density-dependent network structuring within and across wild animal systems

**DOI:** 10.1101/2024.06.28.601262

**Authors:** Gregory F Albery, Daniel J Becker, Josh A Firth, Delphine De Moor, Sanjana Ravindran, Matthew Silk, Amy R Sweeny, Eric Vander Wal, Quinn Webber, Bryony Allen, Simon A Babayan, Sahas Barve, Mike Begon, Richard J Birtles, Theadora A Block, Barbara A Block, Janette E Bradley, Sarah Budischak, Christina Buesching, Sarah J Burthe, Aaron B Carlisle, Jennifer E Caselle, Ciro Cattuto, Alexis S Chaine, Taylor K Chapple, Barbara J Cheney, Timothy Clutton-Brock, Melissa Collier, David J Curnick, Richard J Delahay, Damien R Farine, Andy Fenton, Francesco Ferretti, Laura Feyrer, Helen Fielding, Vivienne Foroughirad, Celine Frere, Michael G Gardner, Eli Geffen, Stephanie S Godfrey, Andrea L Graham, Phil S Hammond, Maik Henrich, Marco Heurich, Paul Hopwood, Amiyaal Ilany, Joseph A Jackson, Nicola Jackson, David MP Jacoby, Ann-Marie Jacoby, Miloš Ježek, Lucinda Kirkpatrick, Alisa Klamm, James A Klarevas-Irby, Sarah Knowles, Lee Koren, Ewa Krzyszczyk, Jillian M Kusch, Xavier Lambin, Jeffrey E Lane, Herwig Leirs, Stephan T Leu, Bruce E Lyon, David W MacDonald, Anastasia E Madsen, Janet Mann, Marta Manser, Joachim Mariën, Apia Massawe, Robbie A McDonald, Kevin Morelle, Johann Mourier, Chris Newman, Kenneth Nussear, Brendah Nyaguthii, Mina Ogino, Laura Ozella, Craig Packer, Yannis P Papastamatiou, Steve Paterson, Eric Payne, Amy B Pedersen, Josephine M Pemberton, Noa Pinter-Wollman, Serge Planes, Aura Raulo, Rolando Rodríguez-Muñoz, Lauren Rudd, Christopher Sabuni, Pratha Sah, Robert J Schallert, Ben C Sheldon, Daizaburo Shizuka, Andrew Sih, David L Sinn, Vincent Sluydts, Orr Spiegel, Sandra Telfer, Courtney A Thomason, David M Tickler, Tom Tregenza, Kimberley VanderWaal, Sam Walmsley, Eric L Walters, Klara M Wanelik, Hal Whitehead, Elodie Wielgus, Jared Wilson-Aggarwal, Caroline Wohlfeil, Shweta Bansal

## Abstract

High population density should drive individuals to more frequently share space and interact, producing better-connected spatial and social networks [1–4]. Although this theory is fundamental to our understanding of disease dynamics [2,5–8], it remains unconfirmed how local density generally drives individuals’ positions within their networks, which reduces our ability to understand and predict density-dependent processes [4,9,10]. Here we provide the first general evidence that density drives greater network connectedness at fine spatiotemporal scales, at the scale of individuals within wild animal populations. We analysed 36 datasets of simultaneous spatial and social behaviour in >58,000 individual animals, spanning 30 species of fish, reptiles, birds, mammals, and insects. 80% of systems exhibited strong positive relationships between local density and network centrality. However, >80% of relationships were nonlinear and 75% became shallower at higher values, signifying that demographic and behavioural processes counteract density’s effects, thereby producing saturating trends [11–15]. Density’s effect was much stronger and less saturating for spatial than social networks, such that individuals become disproportionately spatially connected rather than socially at higher densities. Consequently, ecological processes that depend on spatial connections (e.g. indirect pathogen transmission, resource competition, and territory formation) are likely more density-dependent than those involving social interactions (e.g. direct pathogen transmission, aggression, and social learning). These findings reveal fundamental ecological rules governing societal structuring, with widespread implications. Identifying scaling rules based on processes that generalise across systems, such as these patterns of density dependence, might provide the ability to predict network structures in novel systems.

## Introduction

The number of individuals occupying a given space – i.e., population density – is a central factor governing social systems. At higher densities, individuals are expected to more frequently share space, associate, and interact, producing more-connected spatial and social networks and thereby influencing downstream processes such as mating, learning, and competition. In particular, density-driven increases in network connectedness should provide more opportunities for parasites [1–4,9] or information [16] to spread between hosts[1–4,9] Despite the fundamental nature of such density-dependent processes, evidence is relatively limited that individuals inhabiting higher-density areas have more spatial and social connections. Furthermore, density effects should differ for asynchronous space sharing (e.g. home range overlap) *versus* social associations (e.g. den sharing or grouping) or interactions (e.g. mating or fighting). While several studies have compared animal populations at different densities to demonstrate variation in social association rates among populations (e.g., [15,17,18]) or groups (e.g., [11–13]), attempts to identify such density effects *within* continuous populations of individuals are rarer (but see [14,15,19–21]), and their findings have never been synthesised or compared for spatial and social behaviours. We therefore have an incomplete understanding of how density, as a fundamental ecological parameter, determines socio-spatial dynamics within and across systems. This inhibits our ability to identify and predict how changes in density – e.g. through culling, natural mortality, dispersal, or population booms – influence downstream processes that depend on shared space and social interactions.

The rate at which an individual interacts with conspecifics depends on its spatial and social behaviour within the context of the surrounding environment and population. Adding more individuals into the same space should cause them to more frequently spatially overlap and socially associate or interact (Figure 1). Often, individuals are modelled as randomly moving and interacting molecules (“mass action” or “mean field”). In this conceptualisation, direct contact between two molecules is analogous to a social interaction or association; rates of such interactions are often assumed to increase with density (“density-dependent”; e.g., [22]), and/or to be homogenous in space (e.g., [13]). In reality individuals are unlikely to behave and interact randomly in space, and instead will be influenced by spatially varying factors including local density [10] and competition for resources [15]. Changes in density may cause individuals to alter their foraging behaviour [23–25], dispersal [26,27], social preference or avoidance [14,28], mating behaviour [29], or preferred group size [18]. In some cases, density may have no effect on interaction rates, because individual animals alter their behaviour in a density-dependent manner to maintain a desired interaction rate [30]. These and related processes might produce strong nonlinearities in density-interaction relationships, which can complicate the predictions of density dependence models of pathogen transmission, for example [2,3,9]. For example, individuals or groups can learn to avoid where competitors might go, resulting in greater spatial partitioning under higher densities [31]. Nevertheless, nonlinearities such as these are poorly understood and rarely considered.

**Figure 1:**
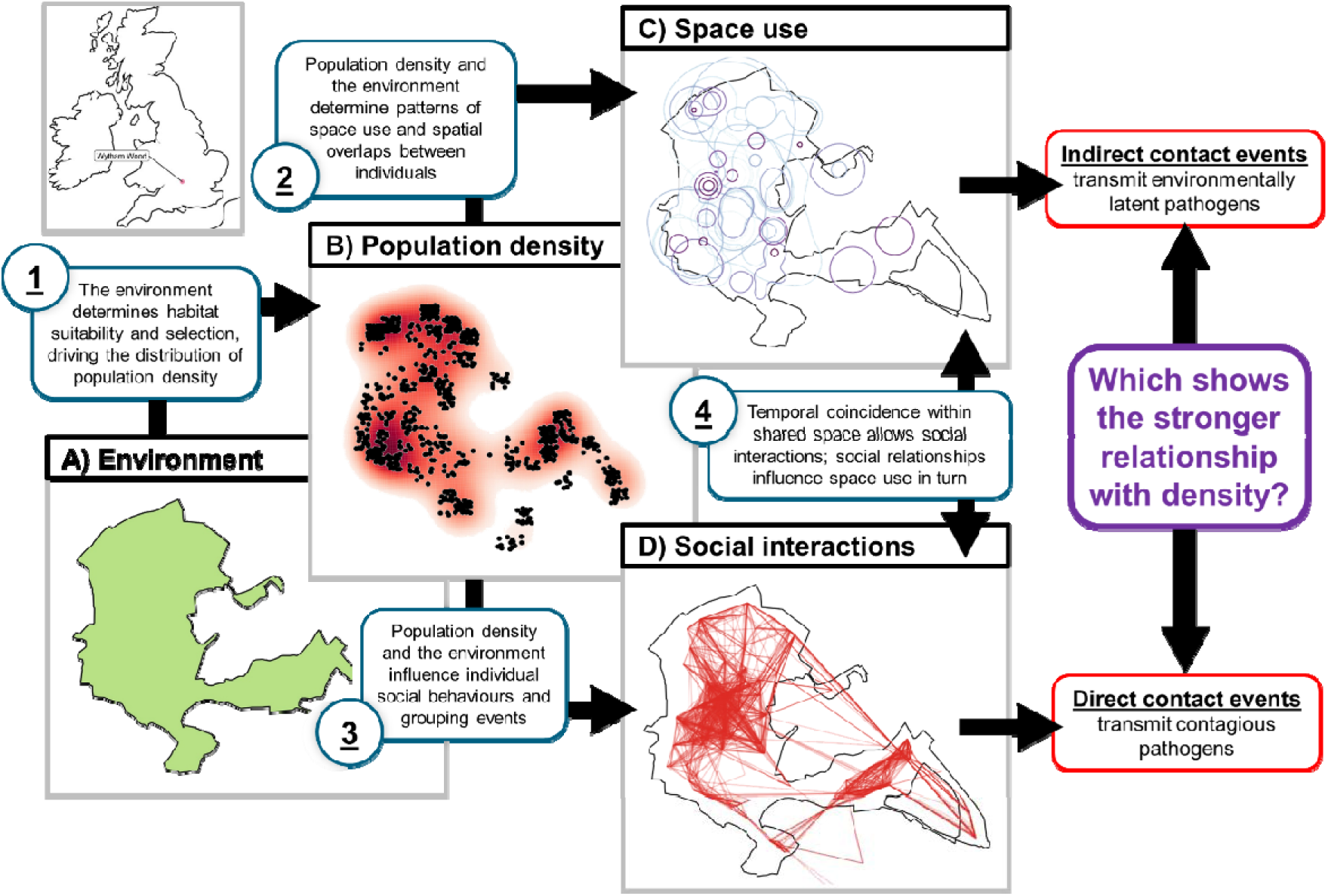
Schematic detailing the rationale underlying this study, outlining how population density drives the formation of spatial and social networks. This depiction uses the Wytham Wood great tits as an example. In panel C, the different purple shades correspond to different individuals’ home ranges. In panel D, red lines depict connections among individuals, with each individual located at their centroid. Ultimately, one of our main aims is to ask whether spatial or social connections generally show a stronger relationship with density, partly functioning as a proxy for indirect and direct contact events with the potential to transmit pathogens. This framework moves between concepts of network and contact formation traversing behavioural ecology, spatial and social network ecology, and disease ecology.

Several wild animal studies have suggested relationships between density and social association rates are often nonlinear and saturating [11–15]. Such relationships imply that association rates do not increase passively with density, but rather that behavioural or demographic processes likely change as density increases, with the ultimate consequence of slowing association rates. However, these nonlinearities are difficult to examine between populations or between species because they introduce a range of confounders and have few replicates along the density axis [9]. On the other end, lower densities may provide less ability to exert social preferences, but low-density populations may be harder to study due to (for example) low return on sampling investment; alternatively, the failure to achieve sufficient interaction rates may result in Allee effects and ultimately drive populations toward decline [32,33].

Characterising gradients of density across individuals within a population offers a workaround to these problems, and facilitates an appreciation of the fact that interactions occur between individuals rather than at the population level. Examining between-individual variation is one reason that social network analysis – which allows characterisation and analysis of individual-level social traits, amongst other things – has become so popular in animal ecology in recent years [34–38]. Additionally, recent years have seen a substantial growth in understanding of socio-spatial behaviours, including harmonising the concepts of spatial and social density [9,10,39]. Applying network analyses coupled with this socio-spatial understanding of density could provide an individual-level picture of density’s effects on spatial and social connectedness, offering far higher resolution and statistical power and greater ability to detect within-system nonlinearities and between-system differences [9]. By providing new understanding of the correlates and emergent consequences of variation in density, this expansion could help to identify general rules underlying social structuring and network scaling in space.

Critically, different types of interactions or associations should show different relationships with density: for example, the need to compete for food at higher densities could drive a disproportionate increase in aggression [40], but this is unlikely to be true of mating interactions. In contrast, higher density and food scarcity should lead to lower exclusivity in resources and more overlapping home ranges, thus enhancing the effect of density on spatial network [41]. This rationale is well-understood in disease ecology, as differences in density-contact relationships are thought to drive differences in density dependence of infection – where “contact” is defined as an interaction or association that could spread a pathogen (Figure 1). “Contacts” then form the basis of spatial and social networks used to investigate pathogen transmission dynamics, which should likewise diverge with density just as contacts do. For example, density should drive greater transmission of respiratory pathogens but not sexually transmitted pathogens [1,42]. Establishing these density-contact relationships is integral to understanding disease dynamics and developing control measures [1,43], but we still have a poor understanding of how different interactions (and therefore contact events for different pathogens) are driven by density. This direct/indirect interaction dichotomy is most fundamental to disease ecology [39,44], but given building interest in the spatial-social interface and relationships between spatial and social networks in behavioural ecology [10], the framework is readily related to other fields (e.g. direct versus indirect cues that can lead to social learning [45]). Previously established density-interaction relationships are diverse and include feral dog bites [19], ant antennations [46] and trophallaxis [30], ungulate group memberships [20,23], rodent co-trapping [11,47], and agamid association patterns [14,21], but no study has yet synthesised how the rates of multiple interaction or association types relate to density, within or across systems.

Identifying the general rules underlying density dependence requires quantifying density’s relationship with proxies of interaction rates at fine scales across a diversity of systems, then identifying the factors determining their slope and shape. To this end, we collate a meta-dataset of over 58,000 individual animals across 36 wildlife systems globally (Figure 2) to ask how within-population variation in density determines between-individual interaction rates based on connectedness in spatial and social networks. We fit multiple competing linear and nonlinear relationships to identify the slope and shape of density effects within each system, and we use meta-analyses to investigate general rules determining their slope and shape across systems. In particular, we focus on comparing space sharing with social interactions and associations as a cross-system case study. Ultimately, we present a *de novo* cross-system analysis of individuals’ social and spatial behaviour that traverses fields of behavioural, population, and disease ecology, which could help to inform general rules governing the structure of social systems, and eventually shape management and conservation decisions in a wide range of systems.

**Figure 2:**
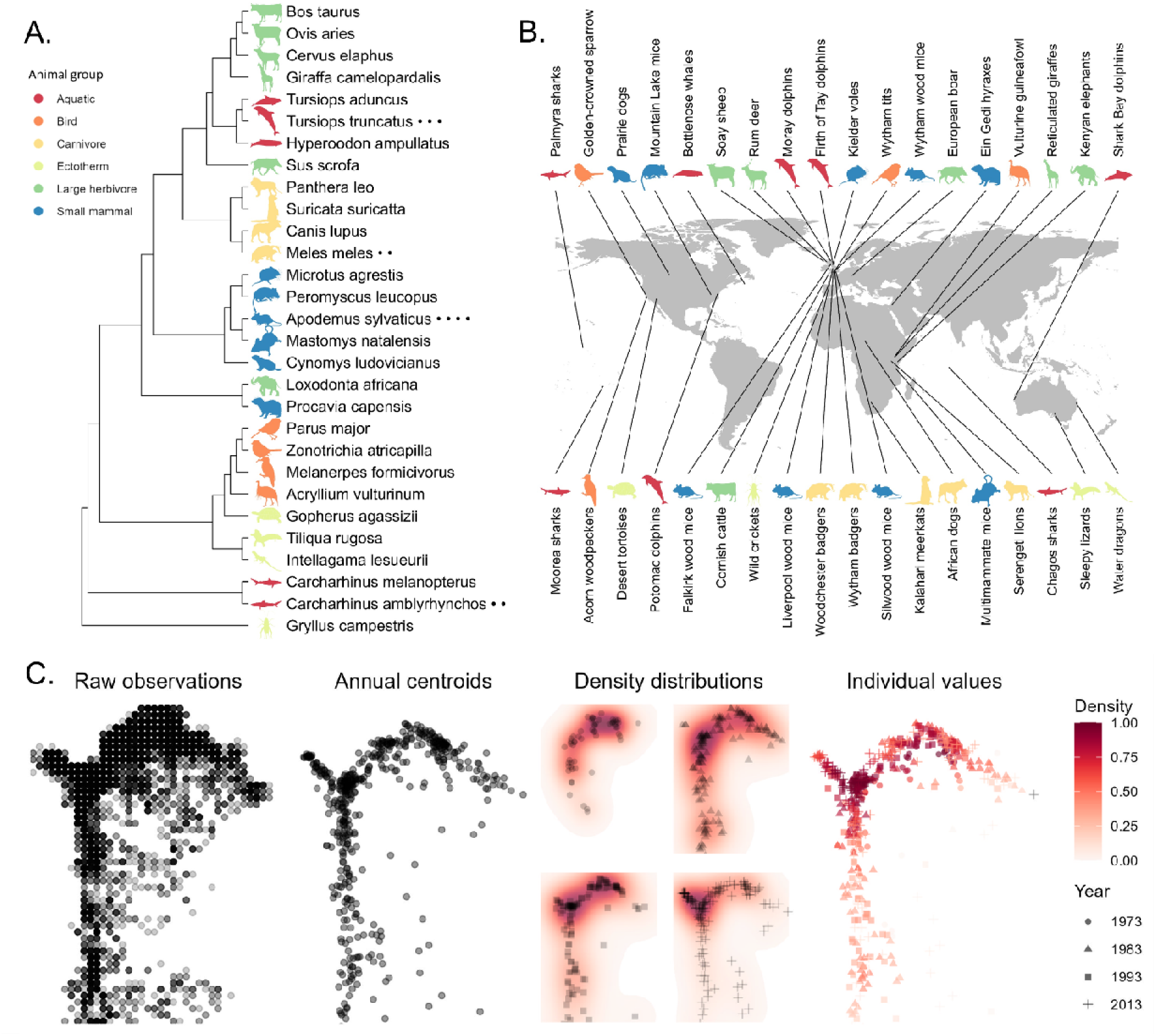
The phylogenetic (A) and geographic (B) distribution of our 36 examined datasets of spatial and social behaviour, with (C) schematic depicting the methodology for deriving local density values, using the Isle of Rum red deer data as an example. The X and Y axes are bivariate spatial coordinates. The panels within (C) show raw observations of individuals in space that we then average at the individual level to make centroids; we use the centroids to generate annual density distributions, which are then assigned to individuals in the form of local density measures. Animal silhouettes are from phylopic.org; a list of attributions is in the supplement (Supplementary Table 2). NB the Potomac dolphins are now defined as Tursiops erebennus; they are currently incorporated in Panel A as T. truncatus, following the Open Tree of Life nomenclature.

## Results and Discussion

We compiled a comparative meta-dataset of over ten million observations of individual animals’ spatial and social behaviour, across a wide range of ecological systems and taxonomic groups of animals. We then ran a standardised pipeline to align their spatial and social observations, identifying strong and predictable relationships between local density and network connectedness at the individual level.

We observed strong positive relationships between individuals’ local population density and their connectedness in spatial and social networks across a wide range of wild animals: of our 64 replicates, 51 (78%) were significantly positive when analysed using linear models (Figure 3A). Meta-analyses identified a highly significant positive mean correlation between density and connectedness, both for social networks (Estimate 0.22; 95% CI 0.17, 0.27) and spatial networks (0.45; 0.36, 0.53; Figure 3B). Our study therefore provides fundamental evidence that high local population density broadly drives greater connectedness within ecological systems, at the individual level. Slopes were highly variable across systems for both spatial and social networks (Figure 3A; Q-test of heterogeneity across systems: Q_37_ = 5627.33 and Q_25_ = 1281.83, both P<0.0001), indicating that quantifying these slopes within and between multiple systems and comparing them is important for understanding animal socio-spatial structure. That is, relationships between density and individual connectedness differ substantially between populations, and the biological mechanisms underlying these divergent trends are likely important. As well as adding resolution and allowing comparisons of density effects across systems, our methodology facilitated fitting of nonlinear relationships (using generalised additive models (GAMs); see below). This approach has only rarely been applied before, and then at much coarser resolution (see [11,12,19]). As such, this study fills an important empirical gap by providing insights into the slope and shape of density-connectedness relationships for a diverse variety of animal groups and their social and spatial behaviours (Figure 4). Nevertheless, despite this diversity, we were able to identify several further general trends in our data.

**Figure 3:**
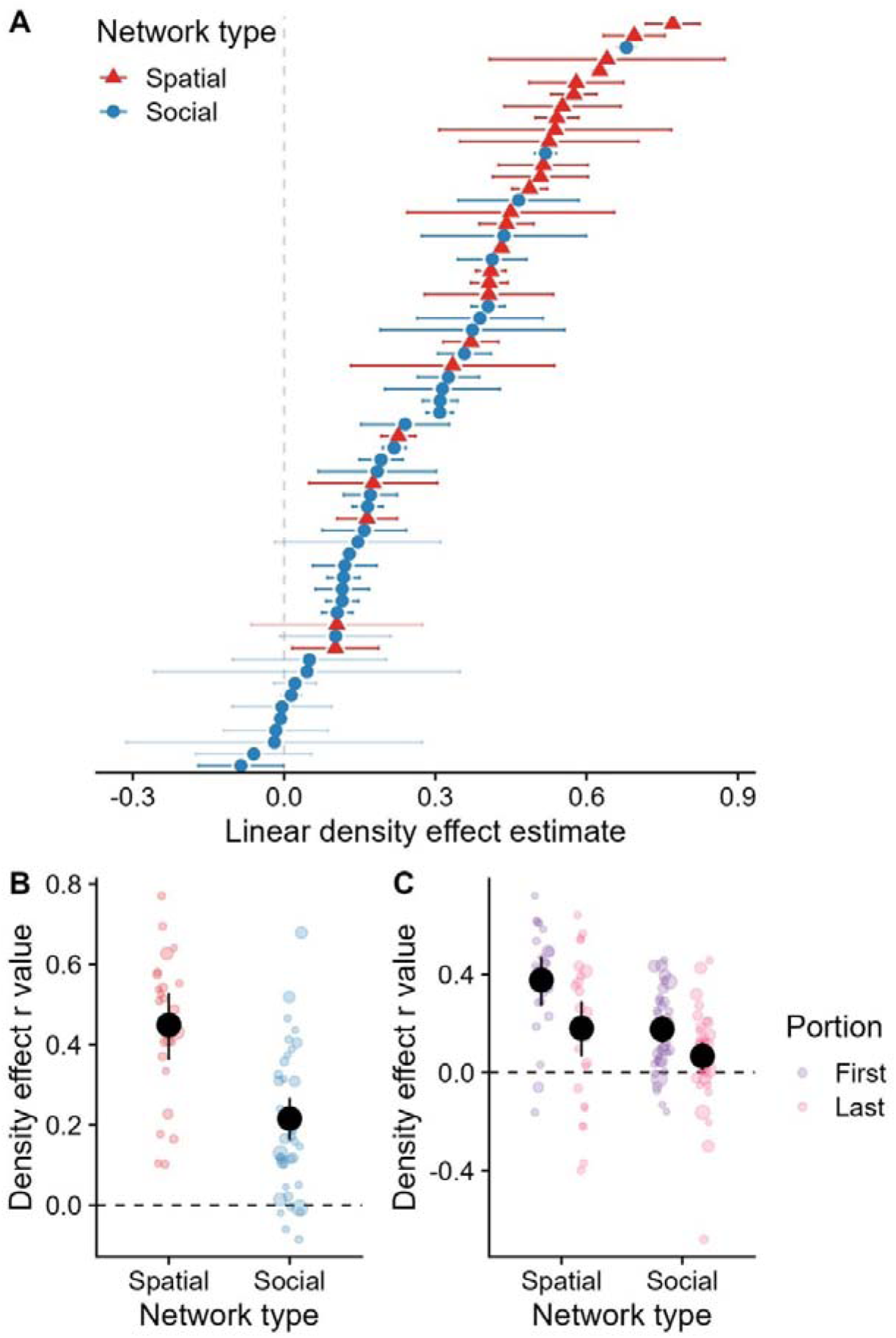
Meta-analysis revealed drivers of variation in linear density effects on individual network connectedness across systems. A) Our fitted linear model estimates of density effects on network strength. Each point represents the mean estimate from a given system; the error bars denote 95% confidence intervals. Opaque error bars were significant (i.e., do not overlap with 0); transparent ones were not. The estimates are in units of standard deviations for both density and network strength. The colour of the point denotes whether the network being examined was defined using spatial or social connections. B) Meta-analyses revealed that centrality in spatial networks (i.e., home range overlap; red points) had a significantly steeper relationship with density than social networks (blue points). C) We fitted linear models separately to two portions of the data within each study population (“first” and “last” represent values below and above the median). The slopes for the latter portion (pink points) were generally less positive than the former portion (purple points), implying a general saturation shape. In panels B) and C), each coloured point represents a study replicate fitted to the strength estimate; points are sized according to sample size, and jittered slightly on the x axis to reduce overplotting. The large black points represent the mean slope estimated from the meta-analysis, and the error bars represent 95% confidence intervals.

**Figure 4:**
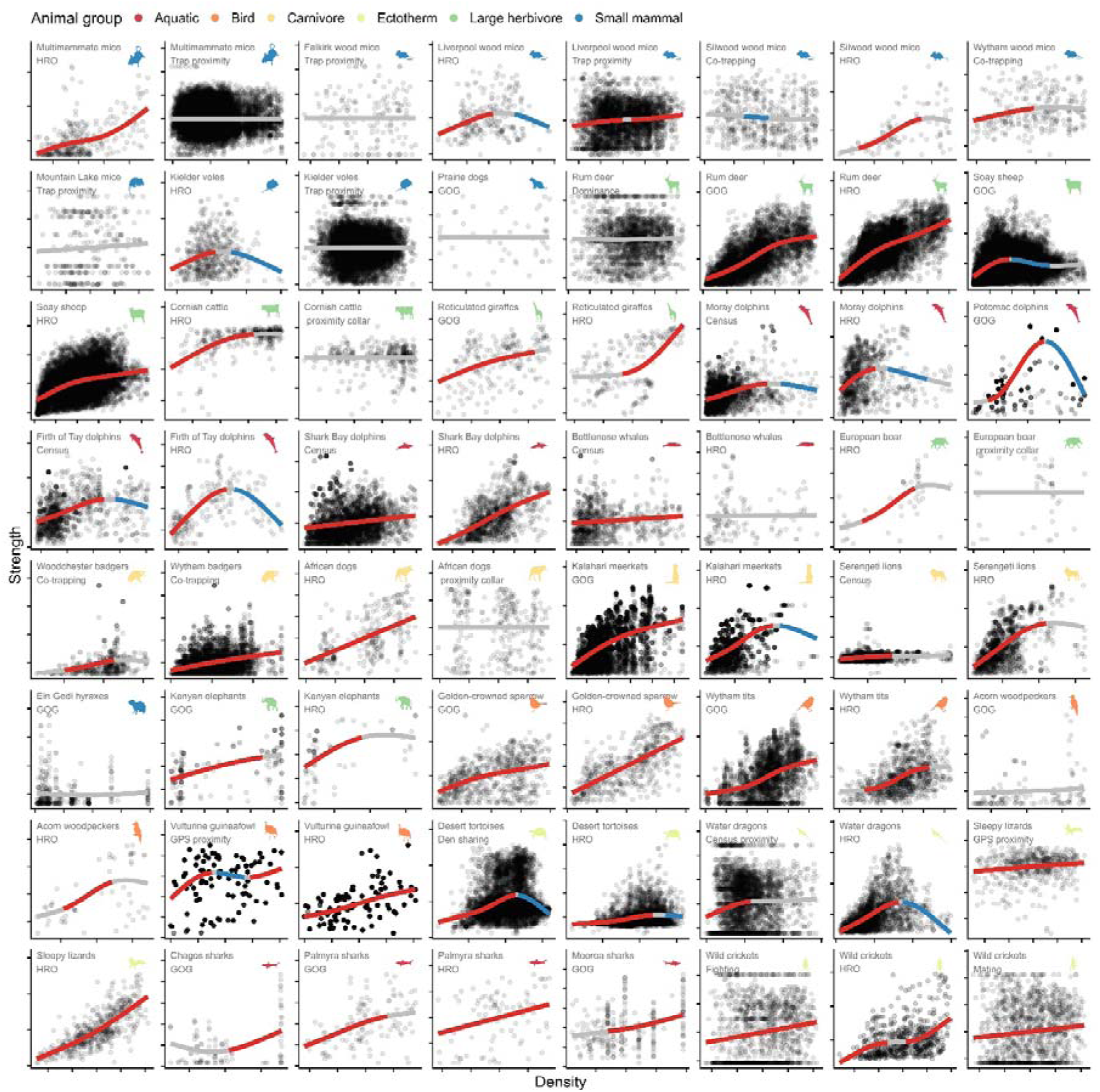
Relationships between density and network connectedness varied substantially across animal systems. Density in individuals per area is on the x axis; network connectedness (strength centrality) is on the y axis. Both values have been standardised to have a mean of zero and a standard deviation of 1 within each system; the axis ticks are in units of 1 standard deviation. Each point represents an individual-year-behaviour replicate; the lines portray the model fit from our generalised additive models (GAMs). Red lengths of the smooth=significantly positive; grey=not significantly different from zero; blue=significantly negative. Points are semi-transparent to enhance visibility. Panels are arranged phylogenetically following the tree displayed in Figure 2A; GOG=gambit of the group; HRO=home range overlap. Animal silhouettes are from phylopic.org; a set of links and attributions are in the Supplement.

Remarkably, density’s effect more than doubled in size for spatial compared to social networks (Figure 3B; r=0.45 *versus* 0.22); there was a difference of 0.26 (CI 0.16, 0.36, P<0.0001) for this effect when we meta-analysed the two contact types together. This finding indicates that as density increases, wild animals are more likely to share space with each other, but that social connections increase at a much slower rate. Similarly, we discovered that saturating shapes were extremely common: as density increased, its effect on connectedness decreased, such that 48/64 systems (75%) had a steeper slope at low density values than at high ones. This effect was strong for both social networks (effect on r= −0.11; CI −0.19, −0.03; P=0.01) and for spatial networks, with substantial overlap between their estimates (−0.22; −0.37, −0.07; P=0.0042). Due to the greater overall effect for space sharing, the latter half of density’s spatial effect was still higher than the first half of its social effect (Figure 3C). Together, these observations suggest that density-dependent processes act to limit the increase in social connectedness with density, but without limiting spatial overlaps to the same extent. Consequently, higher-density areas are characterised disproportionately by individuals asynchronously sharing space rather than socially associating, while in lower-density areas individuals are disproportionately more socially connected proportional to their shared space.

There are many possible social reasons for saturating nonlinearity in density-dependent network structuring: for example, individuals in higher density areas may begin to avoid each other, seeking to avoid competition or aggression [40] or exposure to infectious disease [48]. For instance, Eastern water dragons (*Intellagama lesueurii*) show greater avoidance at higher densities [14], supporting avoidance-related mechanisms. Alternatively, in species with high social cognition or stable bonds, saturation could reflect lower social effort or ability to keep track of social affiliates at higher densities [49]. In general, individuals likely have a preferred social interaction rate or group size – a preference that they may increasingly exert at higher densities [18]. It remains to be seen how this preference varies among individuals, and whether individuals vary in their preferred social network position given a certain density. Given that individuals vary in their movement and spatial phenotypes [50–52], and social phenotypes [52–54] in ways that should manifest for density-dependent behaviours specifically [10], it seems likely that these slopes could vary between individuals as they do between populations. Future analyses might fit variable density-connectedness slopes among individuals to identify socio-spatial syndromes across systems, as has been done previously in single systems including caribou (*Rangifer tarandus*) [55] and American red squirrels (*Tamiasciurus hudsonicus*) [56]. Additionally, we could dissect the social network and its relationship to the spatial network to identify levels of attraction [57,58] or avoidance [59] and how they depend on density.

We considered that density-dependent changes in spatial behaviours might explain these trends: for example, density could create greater competition over resources and therefore reduce energy to roam (and contact others). Individuals may partition their niches [60], or reduce their territory or home range sizes [56,61,62], potentially driven by years of plentiful resources supporting higher densities alongside smaller home ranges sufficiently providing ones’ resource needs, which could drive lower association rates. However, our findings do not seem to support explanations related to small home ranges, because such explanations should produce an equivalent or stronger reduction in (relative) spatial connectedness. In contrast, we observed that density drove individuals to become spatially connected faster than socially, such that the underlying mechanisms likely involve behaviours and demographic processes that specifically affect social collocation in space and time. Testing the precise underlying mechanism will likely require finer-scale behavioural observations, as described below. Regardless of mechanism, these saturating density-connectedness relationships strongly support the idea that examining density effects at the individual level – rather than between populations – is highly informative. For many systems, “mean field” expectations of homogenous interactions under increasing density likely produce an inaccurate (i.e., inflated) picture of density’s effects. Importantly, our study included many examples of proximity-based social networks – most notably “gambit of the group” measures [63] – but relatively few “direct” interactions such as mating, grooming, or fighting. It is interesting that these differences manifested even among two ostensibly spatially-defined contact metrics (gambit of the group and home range overlap). This observation supports the assertion that social association metrics defined by spatiotemporal proximity are valuable for informing on social processes separately from more spatial behaviours *sensu stricto* such as ranging behaviour [20]; we expect that “more direct” interactions could show even further differences in relationships with density. Incorporating a larger number of “direct” metric-based systems could help to address this question (see Supplementary Discussion).

The fact that spatial networks show stronger and more linear density dependence than social networks could heavily influence the ecology of animal systems. For example, indirectly transmitted (i.e., environmentally latent) parasites may exhibit greater density dependence than directly transmitted ones, given that individuals likely experience disproportionately more indirect contact at higher densities. This observation contrasts with orthodoxy that directly transmitted parasites are most likely to be density dependent [64], and supports the value of investigating nonlinear changes in socio-spatial behaviour and grouping patterns in response to density when considering density dependence. Saturating density-connectedness functions further have implications for disease modelling and control. Specifically, our findings lend behavioural support to the growing consensus that many diseases are density-dependent at lower densities, but not at higher densities (i.e., that the slope flattens with density) [22,65]. Rather than assuming constant behavioural mixing at higher densities, epidemiological models could benefit from incorporating density-dependent shifts in behaviours and demography that influence direct and indirect interaction frequencies, as previously suggested empirically and by epidemiological theory [22]. These relationships could influence our targets for culling or vaccination coverage [5]. Given that animals at high density seem likely to have a relatively shallow relationship between density and contact rates, reducing population density – for example by culling – might therefore be ineffective at reducing pathogen transmission initially, particularly when considering socially transmitted pathogens, where contact rates are particularly likely to have become saturated (Figure 3C). Similar problems with culling have already been acknowledged in specific systems – e.g. in canine rabies [43,66,67] – but our study implies that shallow nonlinear density-contact trends could be more general than previously thought and could be driven by flexible density-dependent changes in behaviour and demography. Conversely, culling could be disproportionately effective at intermediate densities and identifying the inflection points of the curve might help to design optimal management strategies. Future studies should investigate whether the divergence in spatial and social connectedness with density drives a concurrent divergence in the prevalence of directly and indirectly transmitted parasites, as well as addressing several other biases in our selection of systems (e.g. [68]; see Supplementary Discussion).

Beyond these general trends, we ran generalised additive models (GAMs) that revealed that 52/64 density effects on network connectedness (81%) were significantly nonlinear (ΔAIC>2); these relationships took a wide variety of shapes, representing a range of nonlinear functions that are hard to generalise (Figure 4). Notably, while many GAM smooths were eventually significantly negative (Figure 4), the vast majority of linear models fitted to the second half of the data were positive (Figure 3C); this result is likely an artefact of restricted model fitting, rather than true downturns in connectedness with density. Nonlinearity did not cluster according to connection type definitions, or according to animal group. These observations were largely corroborated by our meta-analytical models, which found no factors influencing the slope and shape of density effects overall (P>0.05; Supplementary Table 3), including no clear phylogenetic signal (ΔAIC=2.71). This observation speaks to the complexity of these relationships within and across systems, while accentuating that simple functional relationships are often likely to be complicated by contravening ecological factors like habitat selection [69,70], group formation [15], parasite avoidance [71], and demographic structuring [72]. While we were unable to identify specific between-system predictors of nonlinearity of density-connectedness relationships, the finding that most such relationships are strongly nonlinear is an important consideration for future work.

Density is a universal factor underlying the dynamics of animal populations, and its linear and nonlinear effects on spatial and social network structure are likely to impact myriad processes in behaviour, ecology, and evolution. Similar to other studies that have reported general scaling patterns in network analysis [73] and in food web ecology [74], the patterns we report strongly suggest that animal systems generally become more connected spatially than socially under increasing density. These trends might extrapolate to human networks, given that other scaling patterns in animal networks do [73]. As these patterns seemingly manifest regardless of animal group and interaction type, they may reflect a generalisable rule governing the socio-spatial structure of ecological systems. Further refining and implementing these models could facilitate prediction of network structure in novel systems.

Finally, this study is relatively unique in conducting an expansive meta-analysis of behavioural data from individual animals across a diverse selection of systems. As datasets accumulate comparative analyses are increasing in frequency in social network ecology [75], but often revolve around analysing whole networks rather than individuals [76], and never (to our knowledge) in conjunction with analyses of spatial behaviour. These analyses therefore hold exceptional promise for disentangling spatial and social behaviour across diverse systems; for example, given that our dataset includes many repeatedly sampled known individuals, future analyses could investigate individual-level repeatability or multi-behaviour “behavioural syndromes” across a variety of different taxa and environments [10,77]. Additionally, capitalising on the wide range of methodological approaches to behavioural data collection (e.g. censuses, trapping, and GPS telemetry), the methodological constraints of socio-spatial analyses could be tested in this wide meta-dataset as they have been in other recent comparative analyses of wild ungulates [78]. As well as being diverse, our meta-dataset had several replicate examples of (for example) marine mammals and trapped rodents, which could be used for finer-scale and more targeted comparative analyses within these smaller taxonomic groupings. For now, it is highly encouraging that we uncovered general trends across these disparate animal systems, and further explorations of these socio-spatial patterns may help to inform a wide range of exciting and longstanding questions at the spatial-social interface [10].

## Methods

### Data standardisation and behavioural pipeline

Data were manipulated and analysed using R version 4.2.3 [79], and all R code is available at https://github.com/gfalbery/DensityMetaAnalysis. Our 36 datasets each involved at least one continuous uninterrupted spatial distribution of observations in a single population; some datasets comprised multiple such populations; all systems had at least one social network measure, and two had two different types of social interaction. These datasets covered 30 different animal species, including sharks, carnivores, cetaceans, ungulates, rodents, elephants, birds, reptiles, and one orthopteran insect (Figure 2). In one case (The Firth of Tay and Moray Dolphins) we used two distinct replicates despite being composed of overlapping groups of individuals, because of their distinct spatial distributions, which made it difficult to fit a coherent density distribution.

To standardise the timescale across studies, all systems were analysed as annual replicates – i.e., social and spatial networks were summarised within each year. Our analyses used 64 system-behaviour replicates, listed in Supplementary Table 1, and totalled 151,835 unique system-individual-year-behaviour data points.

All spatial coordinates were converted to the scale of kilometres or metres to allow comparison across systems. To provide an approximation of local density, following prior methodology [20,80], we took each individual’s average location across the year (their centroid) and created a spatial density kernel using the ‘adehabitathr’ package [81], which provides a probabilistic distribution of population density across each study system based on the local frequencies of observed individuals. Each individual was assigned an annual estimate of local density based on their centroid’s location within this spatial density distribution. We made these density distributions as comparable as possible between systems by incorporating the density raster using metre squares; however, there were large differences in density across populations that were difficult to resolve and put on the same scale (e.g. interactions per individual/km^2^ unit of density). Consequently, we scaled and centred density to have a mean of zero and a standard deviation of one within each population, which allowed us to focus on differences in relative slope and shape across systems.

To validate the local density measures estimated using the kernel density approach, we also estimated local density for individuals across all populations based on the locations of individual annual centroids within a designated area. To do so, we first estimated the area of the minimum bounding box (MBB) within which all individuals were censused during the study period based on their annual centroids. For each individuals mean location, we then estimated a circular boundary of radius r=1/20 * area of MBB. We then calculated the number of individuals present within this boundary as an individual’s local density measure. We estimated the Pearson correlation coefficients between the local density measures derived using the KDE approach and the proportional area - based approach (Supplementary Figure 1).

To provide a measure of asynchronous space sharing, we constructed home range overlap (HRO) networks based on proportional overlap of two individuals’ minimum convex polygon (MCP; i.e., the bounding polygon around all observations of each individual in a given year). These HRO networks were restricted to only individuals with five or more observations in a given year to allow us to create convex polygons effectively; 10/36 (28%) systems did not have sufficient sampling for this analysis. We also repeated our analyses with a series of higher sampling requirements for observation numbers to ensure that our findings were robust to this assumption. The MCP approach is relatively low-resolution, and assumes uniform space use across an individual’s home range; however, this approach is less data intensive – and less sensitive to assumptions – than density kernel-based approaches that would estimate variation in space use across the home range, allowing us to apply the models across more systems, more generalisably, and more conservatively.

To provide a measure of social connectedness, we built social networks using various approaches as defined by the original studies: direct observations of dyadic interactions (e.g. fighting or mating); gambit of the group (GoG; i.e., membership of the same group) [63]; co-trapping (i.e., trapped together or in adjacent traps within a given number of trapping sessions); or direct contact measured by proximity sensors (defined by a certain distance-based detection threshold). Notably some analyses use indirect interactions – i.e., spatial overlap – to *approximate* direct interactions, which requires spatiotemporal coincidence, which we caution against particularly when modelling pathogen transmission [39,82]. While the two do often correlate, here we are not using HRO to approximate direct interaction rates, but rather as a measure of indirect interactions (e.g., indicative of transmission of environmental parasites).

For each social network, we scaled connection strength relative to the number of observations of each individual in a dyad (i.e., simple ratio index [83]). Our response variable therefore took the form of strength centrality, scaled to between 0-1 for each dyad, for each social and spatial network. We focus on comparing density effects on social interactions and associations with density’s effects on space sharing.

### Density-connectedness models: what forms do density effects take?

We developed a novel workflow to allow us to derive and compare density’s effects on connectedness – and their drivers – in a standardised way across our animal systems. We fitted models with three main forms: **linear models** fitted to the whole dataset, nonlinear **Generalised additive models** fitted to the whole dataset, and linear **saturation models** fitted separately to low- and high-density subsets of each dataset.

**Linear models:** For each system-behaviour replicate, we first fitted a linear model using the ‘lm’ function in R, fitting scaled density as an explanatory variable to estimate linear density effect slopes. The linear fits are displayed in the supplement (Supplementary Figure 2), as are the residuals (Supplementary Figure 3).

**Generalised additive models (GAMs):** We fitted GAMs in the ‘mgcv’ package [84] to identify whether each density effect was better described by a linear or nonlinear relationship, and to identify the shape of these nonlinear relationships. For each model, we fitted a default thin plate spline with k=4 knots. This knot number was selected to reduce overfitting in our models, which formed several fits to the data that were difficult to reconcile with functional formats. To assess whether nonlinear models fit better than linear models, we used Akaike Information Criterion (AIC), with a contrast of 2ΔAIC designated to distinguish between models.

**Saturation models:** To quantify whether density effects were generally saturating (i.e., that density had steeper relationships with individuals’ connectedness at lower density values), we split the data into two portions: all values below the median density value, and all values above the median. We then re-ran linear models examining the relationship between density and strength in each portion. We attempted to investigate nonlinear patterns (especially saturating effects) across all our systems using a range of other methods (e.g., comparing specific functional relationships with nonlinear least squares), but found that they were generally incapable of fitting well to the data in a standardised way across the many datasets (i.e., non-convergence of nonlinear least squares using semi-automated starting estimates across systems). As such, this approach represented a tradeoff between tractable, generalisable model fitting, interpretability, and accurate representation of the relationship’s shape. All else being equal, we posit that investigating the relative slopes of two otherwise-identical portions of the data is a conservative and informative method of identifying saturation, which was our main hypothesis for the expected shape of density effects.

**Heteroskedasticity and log-log models:** To ensure that our estimates were robust to non-normality and to provide another source of information concerning possible saturation effects, we also conducted tests of heteroskedasticity on our linear models and accompanied them with simulations and fitted log-log linear models. First, we carried out a simple simulation study to test how: a) the skew in residuals; b) a saturating relationship; and c) heteroscedasticity impact whether we may under- or overestimate the slope of an assumed linear relationship between density and strength (See Supplementary Methods - Heteroskedasticity Simulations). These demonstrated that our models were resilient to skew and saturating effects, but that heteroskedasticity in residuals could drive overestimated linear effects in our models.

To examine this possibility further, we derived the Breusch-Pagan statistic for each linear model as a measure of heteroskedasticity, and then plotted it against the meta-analysis covariates and fixed effects. There was no evidence that the density effect was being skewed to be greater for spatial behaviours due to heteroskedasticity, and neither were the second portions of the data more heteroskedastic, which would be expected if this was driving the saturating effect (Supplementary Figure 4). Finally, we fitted log-log linear models with the same formulations as our main linear models defined above, but with both density and strength log(X+1)-transformed, rather than scaled to have a mean of 0 and a standard deviation of 1 (Supplementary Figure. Our results showed broadly identical findings of greater estimates for spatial behaviours, and the fact that the slopes were largely under 1 is indicative of a saturating effect. As such, these tests strongly support our findings’ resilience to uneven data distributions.

### Meta-analysis: what factors determine the slope of density-connectedness relationships?

To characterise the typical relative slope of density effects across systems and identify the factors influencing their variation, we fitted hierarchical meta-analytical models using the ‘metafor’ package in R. The response variable was the standardised slope of the linear density effect; because both individual network strength and density were scaled to have mean of zero and standard deviation of one in the linear regression, this is equivalent to the correlation coefficient (*r*) [85]. We converted all correlation coefficients into Fisher’s *Z* (*Z_r_*) and computed associated sampling variance.

For our hierarchical meta-analysis models, we used an initial model that nested observations within a system-level random effect to account for within- and between-system heterogeneity [86], as 26/36 systems had more than one density effect. We used another random effect for species to account for repeat observations per animal species.

We then added a separate random effect for animal phylogeny [87]. This effect used a phylogenetic correlation matrix of our 30 animal species derived from the Open Tree of Life via the ‘rotl’ package [88], with the ‘apè package used to resolve multichotomies and provide branch lengths [89].

We then fitted intercept-only models using the ‘rma.mv()’ function with restricted maximum likelihood (REML), weighted by inverse sampling variance, and used variance components to quantify *I^2^*, the contribution of true heterogeneity to the total variance in effect size. We used Cochran’s Q to test whether such heterogeneity was greater than that expected by sampling error alone.

We next fitted models with the same random effects structure that included explanatory variables. To detect whether some animals were more likely to experience density effects, we fitted **Animal group** as a factor with six categories, representing a combination of species’ taxonomy and general ecology: aquatic (fish and dolphins), birds, large herbivores (elephants and ungulates), small mammals (rodents and hyraxes), carnivores, and ectotherms (insects and reptiles). We also fitted several explanatory variables indicative of greater statistical power that might increase the strength of density effects: **Geographic area** (km^2^, log_10_-transformed), **Number of years** of study, and **Number of individuals**, all of which we fitted as continuous covariates. Broadly, the animal group model was highly uninformative and competed with the other effects, and we expected that the phylogeny would be more informative, so we report the results of the model without the animal group effect fitted.

We ran several different versions of these meta-analyses: first, we fitted meta-analytical models to the **overall linear models** of spatial and social interaction types separately, and then together, to investigate differences between the spatial and social networks in terms of their mean density slope. Next, we fitted duplicated versions of these models, but with the **saturation models**. These models were identical, but each system replicate had two linear estimates: one taken from the first 50% of the data (up to the median), and one from the latter 50%. By fitting a binary fixed effect of “data portion” to the meta-analytical models, this model would tell us whether the slopes were generally higher in the first portion of the data than the last (and therefore showed generally saturating shapes). We were unable to fit meta-analytical models to our GAMMs, as methods for the meta-analysis of nonlinear estimates are not yet well defined.

## Acknowledgements

GFA and SB were supported by NSF grant number DEB-2211287. GFA acknowledges support from a College for Life Sciences Fellowship at the Wissenschaftskolleg zu Berlin and WAI (C-2023-00057). DJB acknowledges support from the Edward Mallinckrodt, Jr. Foundation and NSF BII 2213854. JAF acknowledges funding from BBSRC (BB/S009752/1), NERC (NE/S010335/1 and NE/V013483/1), and WAI (C-2023-00057). For a list of system-specific acknowledgements, please see the supplement.

## Code availability

All R code is available at https://github.com/gfalbery/DensityMetaAnalysis.

## Supplement

**Supplementary Table 1:**
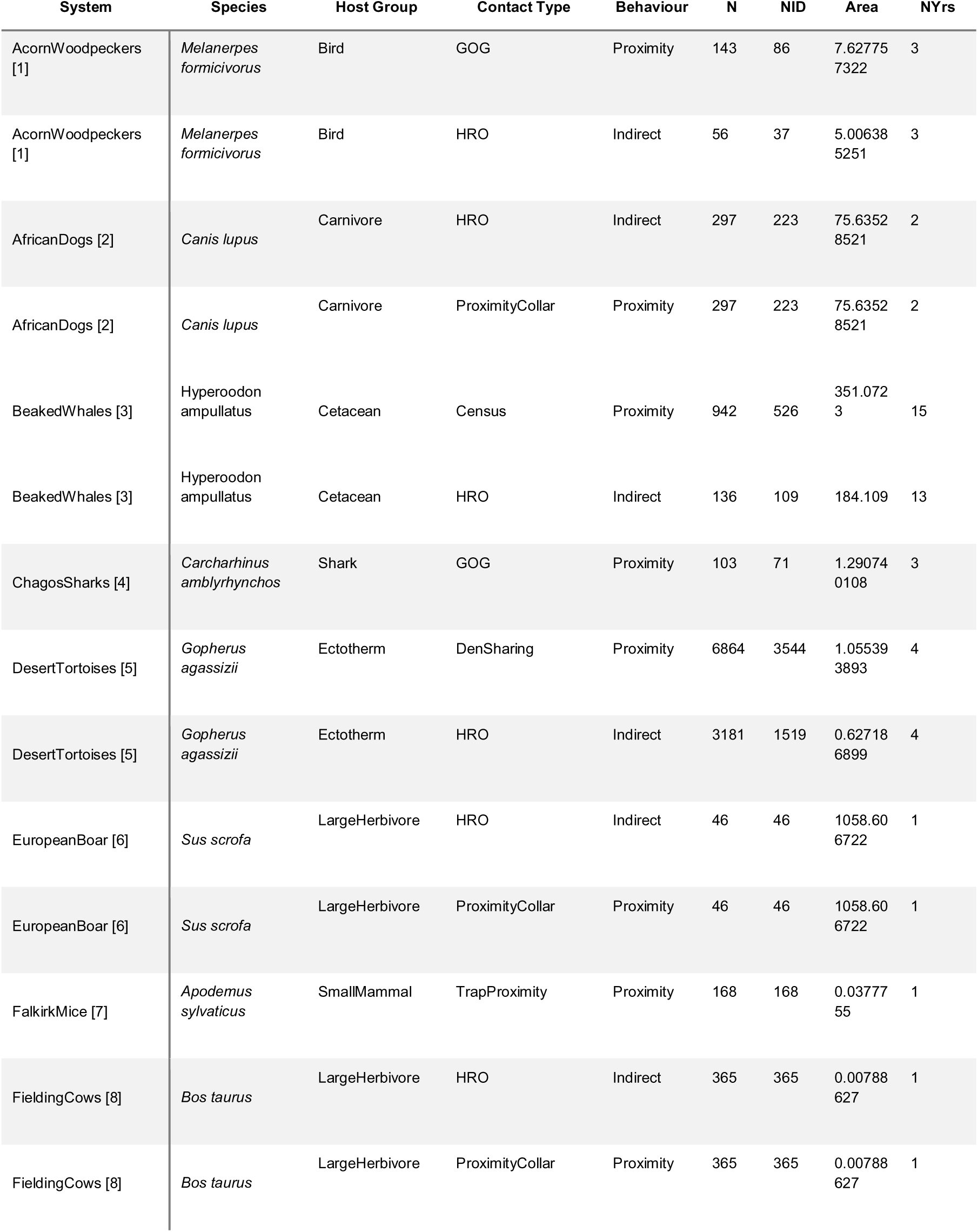
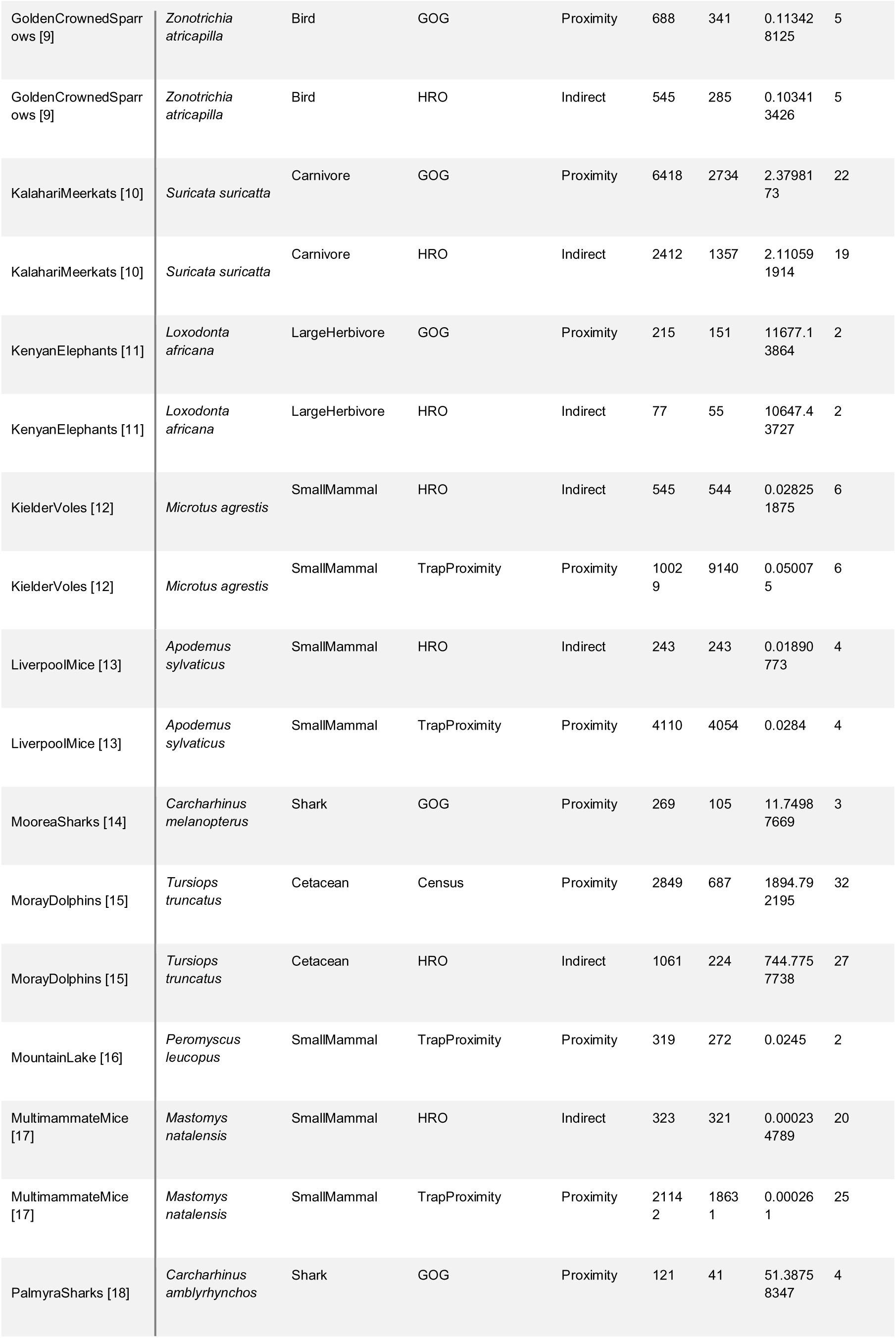
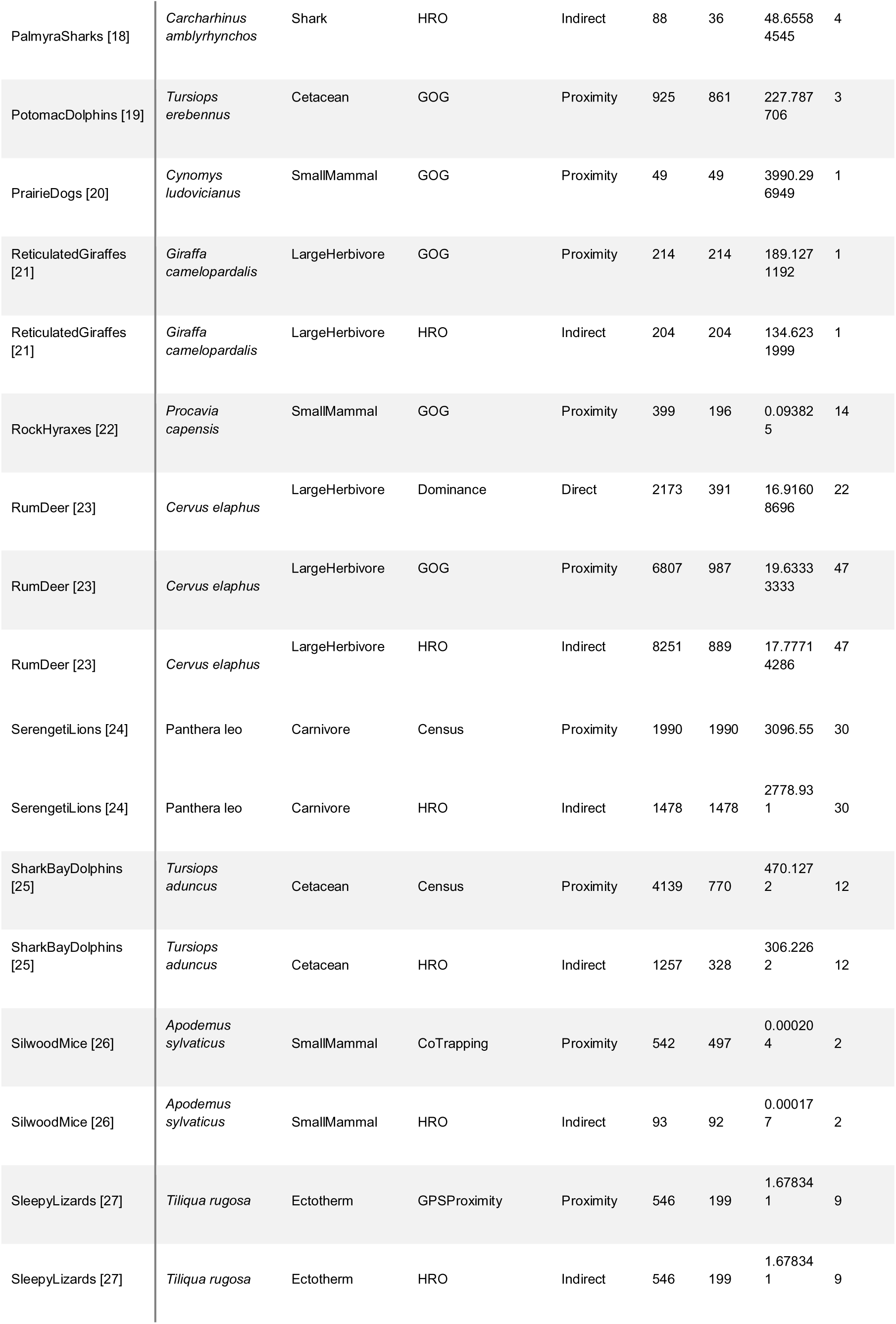
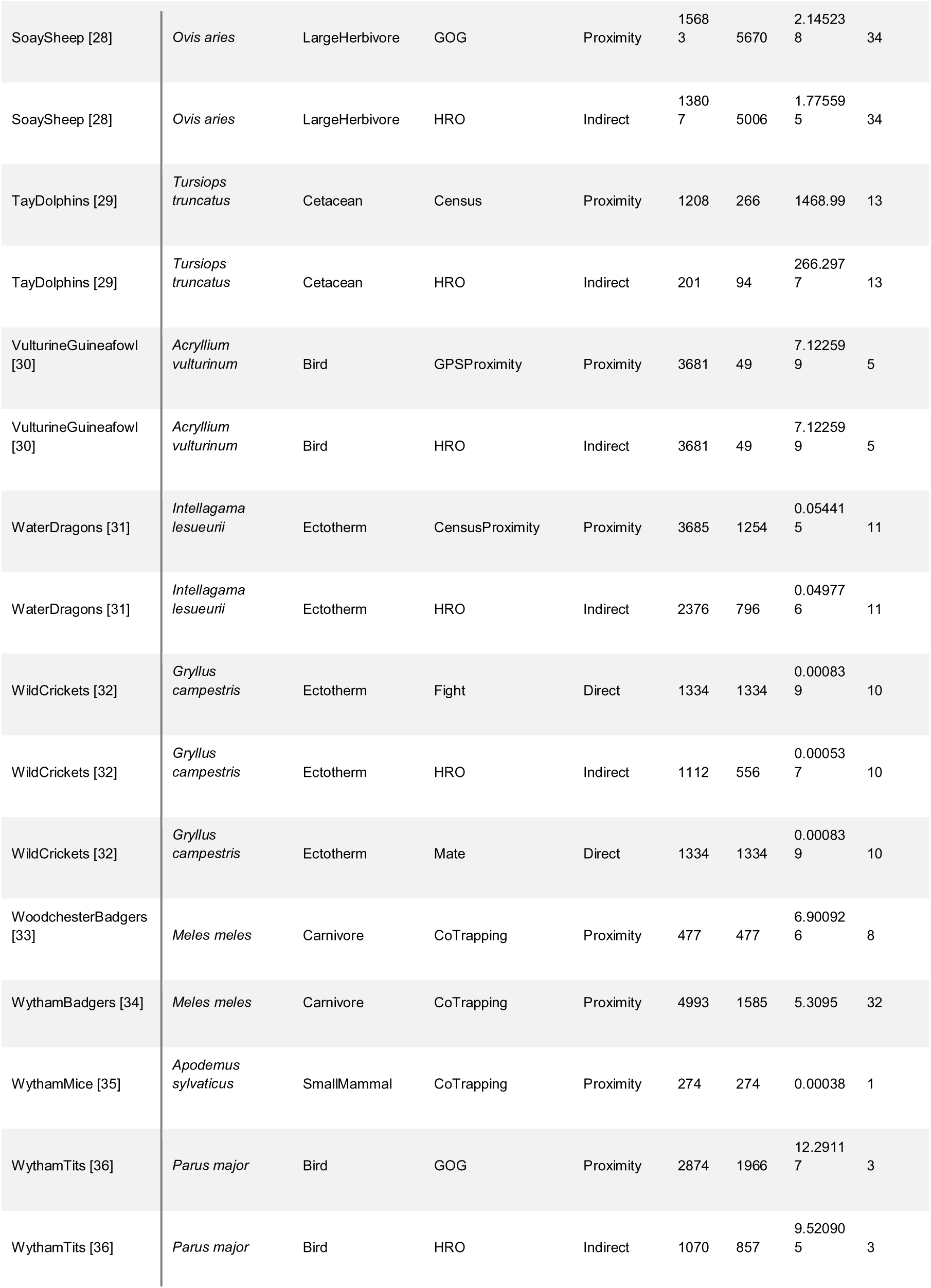
List of study system replicates and their associated traits, with an example reference from each. Also available as a supplementary file. N=number of individual-by-year values; NID=number of unique individuals. Area=study extent in Km^2^. NYears=number of years covered by the study.

**Supplementary Table 2:** The phylopic.org images used in the main text and their associated licenses.

| System | Species | Image credit | License link |
| --- | --- | --- | --- |
| VulturineGuinea fowl | *Acryllium vulturinum* | James Klarevas | https://creativecommons.org/licenses/by/4.0/ |
| WythamMice | *Apodemus sylvaticus* | Anthony Caravaggi | https://creativecommons.org/licenses/by-nc-sa/3.0/ |
| SilwoodMice | *Apodemus sylvaticus* | NA | Public domain |
| LiverpoolMice | *Apodemus sylvaticus* | NA | Public domain |
| FalkirkMice | *Apodemus sylvaticus* | NA | Public domain |
| FieldingCows | *Bos taurus* | NA | Public domain |
| AfricanDogs | *Canis lupus* | NA | Public domain |
| ChagosSharks | *Carcharhinus amblyrhynchos* | NA | Public domain |
| PalmyraSharks | *Carcharhinus amblyrhynchos* | Russell Engelman | https://creativecommons.org/licenses/by/4.0/ |
| MooreaSharks | *Carcharhinus melanopterus* | Ignacio Contreras | https://creativecommons.org/licenses/by/3.0/ |
| RumDeer | *Cervus elaphus* | NA | Public domain |
| PrairieDogs | *Cynomys ludovicianus* | NA | Public domain |
| ReticulatedGiraffes | *Giraffa camelopardalis* | Cathy | https://creativecommons.org/licenses/by-nc-sa/3.0/ |
| DesertTortoises | *Gopherus agassizii* | Andrew A. Farke, shell lines added by Yan Wong | https://creativecommons.org/licenses/by/3.0/ |
| WildCrickets | *Gryllus campestris* | NA | Public domain |
| WaterDragons | *Intellagama lesueurii* | NA | Public domain |
| KenyanElephants | *Loxodonta africana* | Agnello Picorelli | https://creativecommons.org/licenses/by-nc-sa/3.0/ |
| Multimammate Mice | *Mastomys natalensis* | David Liao | https://creativecommons.org/licenses/by-sa/3.0/ |
| AcornWoodpeckers | *Melanerpes formicivorus* | NA | Public domain |
| WoodchesterBadgers | *Meles meles* | NA | Public domain |
| WythamBadgers | *Meles meles* | Anthony Caravaggi | https://creativecommons.org/licenses/by-nc-sa/3.0/ |
| KielderVoles | *Microtus agrestis* | NA | Public domain |
| SoaySheep | *Ovis aries* | Gabriela Palomo-Munoz | https://creativecommons.org/licenses/by-nc/3.0/ |
| WythamTits | *Parus major* | NA | Public domain |
| MountainLake | *Peromyscus leucopus* | Nina Skinner | https://creativecommons.org/licenses/by/3.0/ |
| RockHyraxes | *Procavia capensis* | NA | Public domain |
| KalahariMeerkats | *Suricata suricatta* | NA | Public domain |
| GermanBoar | *Sus scrofa* | NA | Public domain |
| SleepyLizards | *Tiliqua rugosa* | CNZdenek | https://creativecommons.org/licenses/by-nc/3.0/ |
| PotomacDolphins | *Tursiops truncatus* | Chris huh | https://creativecommons.org/licenses/by-sa/3.0/ |
| MorayDolphins | *Tursiops truncatus* | NA | Public domain |
| AberdeenDolphins | *Tursiops truncatus* | Chris huh | https://creativecommons.org/licenses/by-sa/3.0/ |
| SharkBayDolphins | *Tursiops truncatus* | Chris huh | https://creativecommons.org/licenses/by-sa/3.0/ |
| GoldenCrowne<br>dSparrows | *Zonotrichia atricapilla* | NA | Public domain |
| SerengetiLions | *Panthera leo* | NA | Public domain |
| BeakedWhales | *Hyperoodon ampullatus* | Chris huh | https://creativecommons.org/licenses/by-sa/3.0/ |

**Supplementary Table 3:**
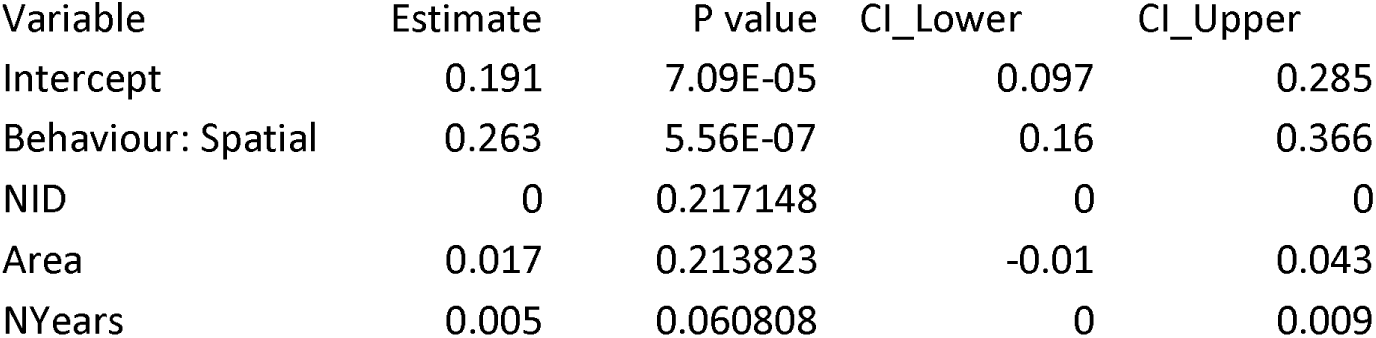
Meta-analysis effect estimates from the full meta-analytical model, providing the estimate, P value, and 95% confidence intervals.

## Supplementary figures

**Supplementary Figure 1:**
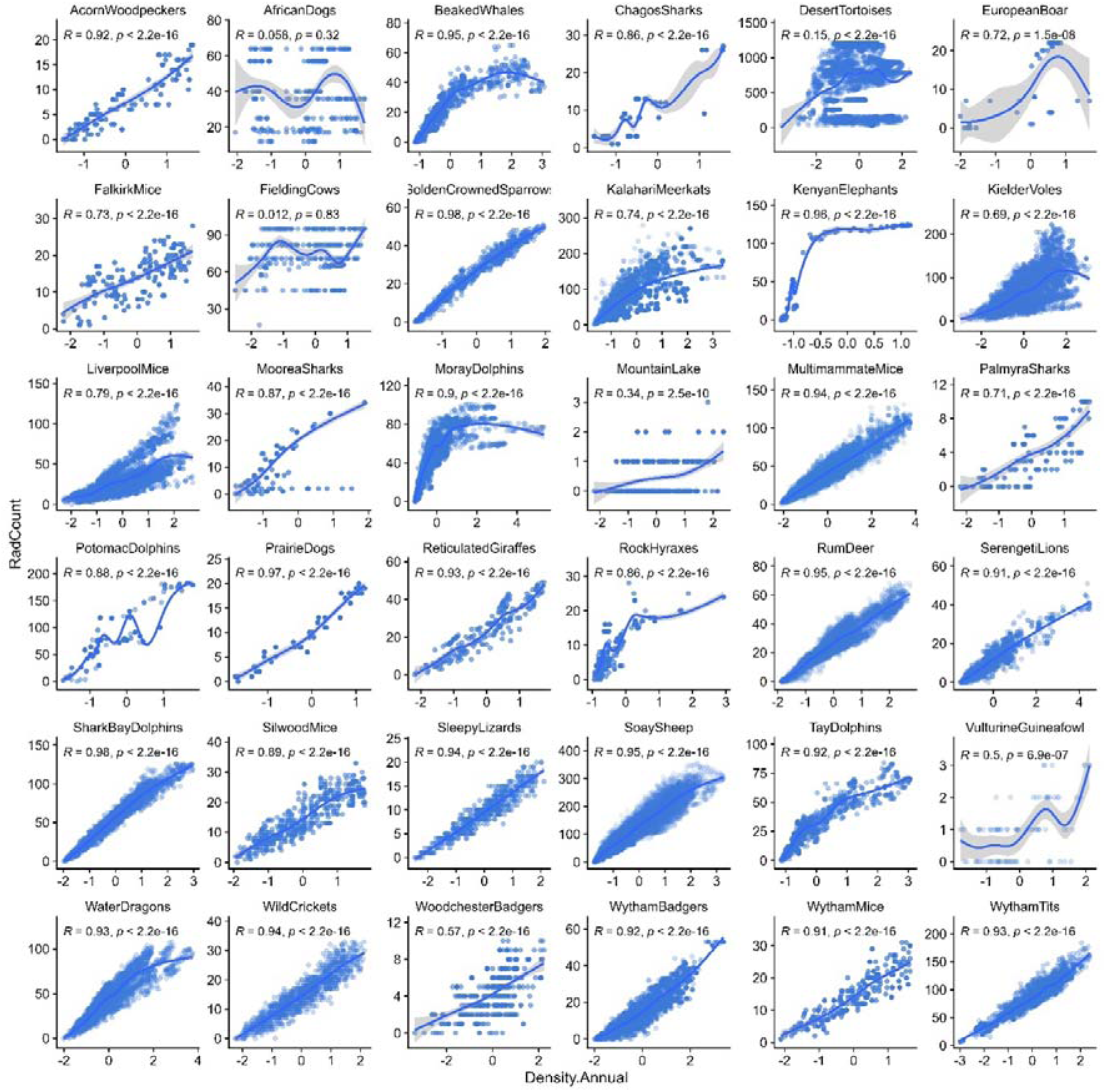
Relationships between two measures of local population density across animal systems. Density on the x axis was calculated using a kernel density estimation approach; density on the y axis was calculated based on the number of individuals located within a radius defined by the study system’s area. X axis values have been standardised to have a mean of zero and a standard deviation of 1 within each site; the axis ticks are in units of 1 standard deviation. Each point represents an individual-year-behaviour replicate; the lines portray the model fit from a generalised additive model (GAM). Correlation coefficients were calculated using a Spearman’s rank correlation.

**Supplementary Figure 2:**
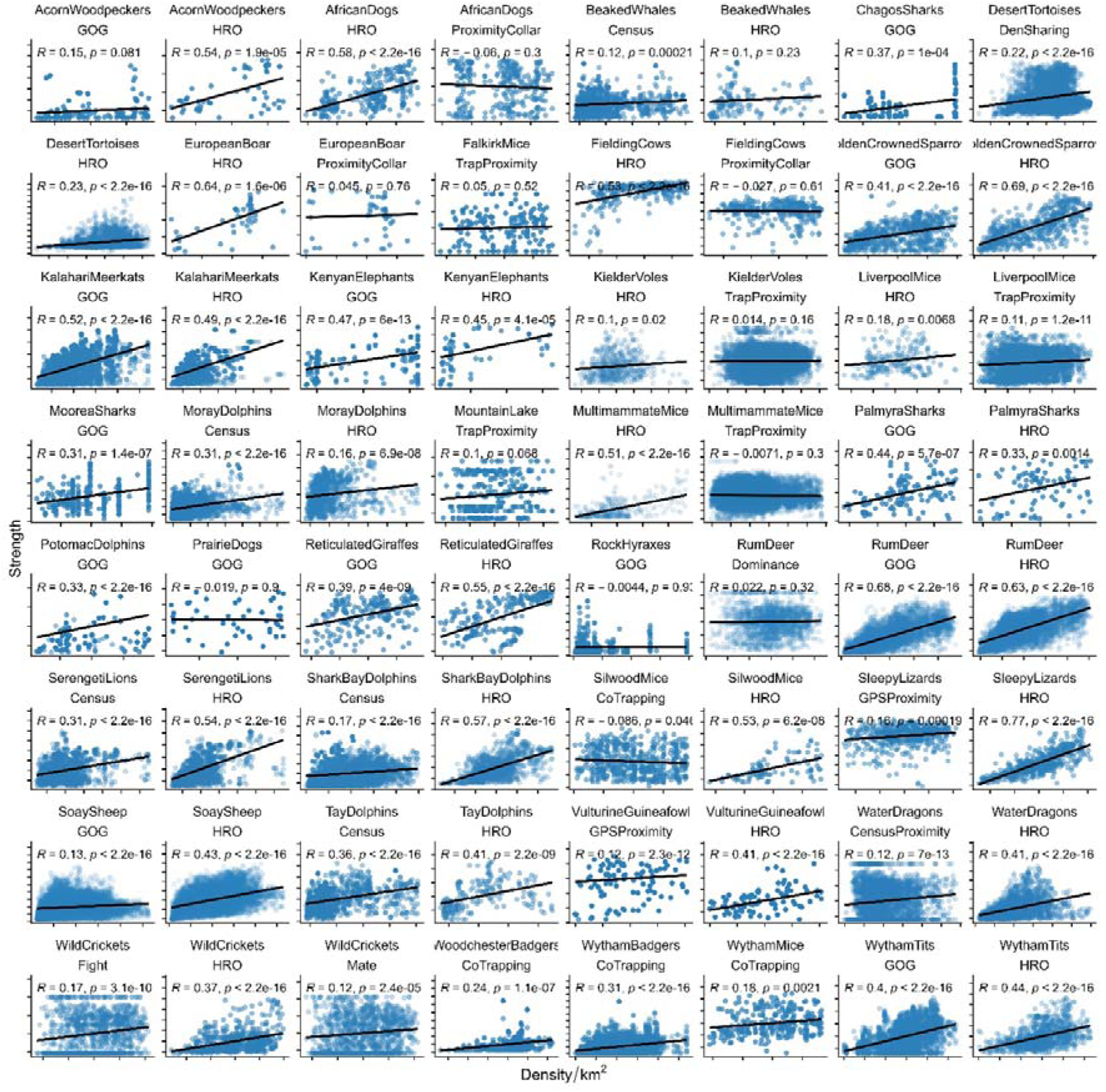
Linear relationships between density and network connectedness across animal systems. Density in individuals per area is on the x axis; network connectedness (strength centrality) is on the y axis. Both values have been standardised to have a mean of zero and a standard deviation of 1 within each system; the axis ticks are in units of 1 standard deviation. Each point represents an individual-year-behaviour replicate; the lines portray the model fit from our linear models for meta-analysis. Points are semi-transparent to enhance visibility. Panels are arranged phylogenetically following the tree displayed in Figure 2A; GOG=gambit of the group; HRO=home range overlap.

**Supplementary Figure 3:**
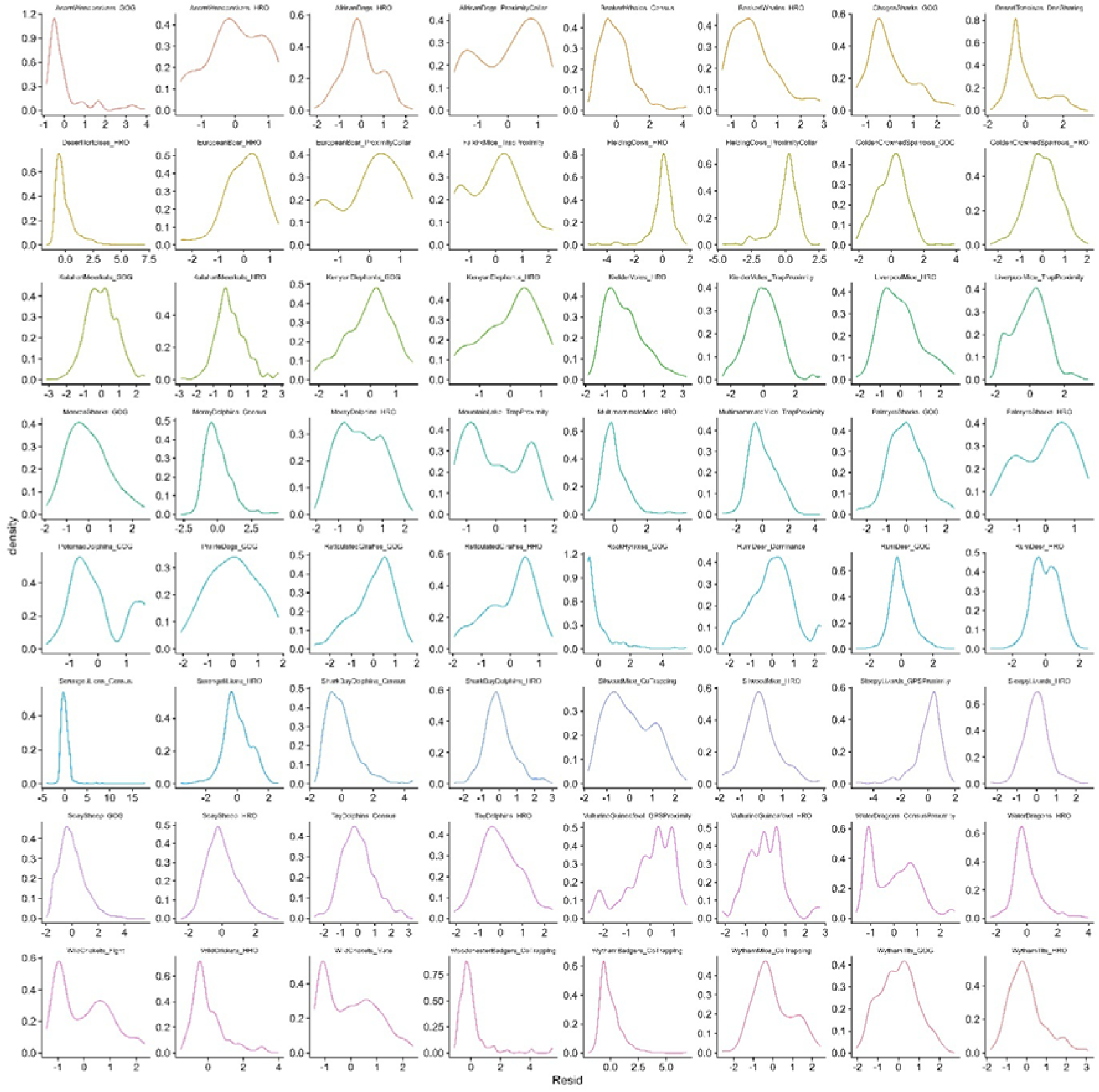
Distributions of residuals from the linear models displayed in Figure 2 above.

**Supplementary Figure 4:**
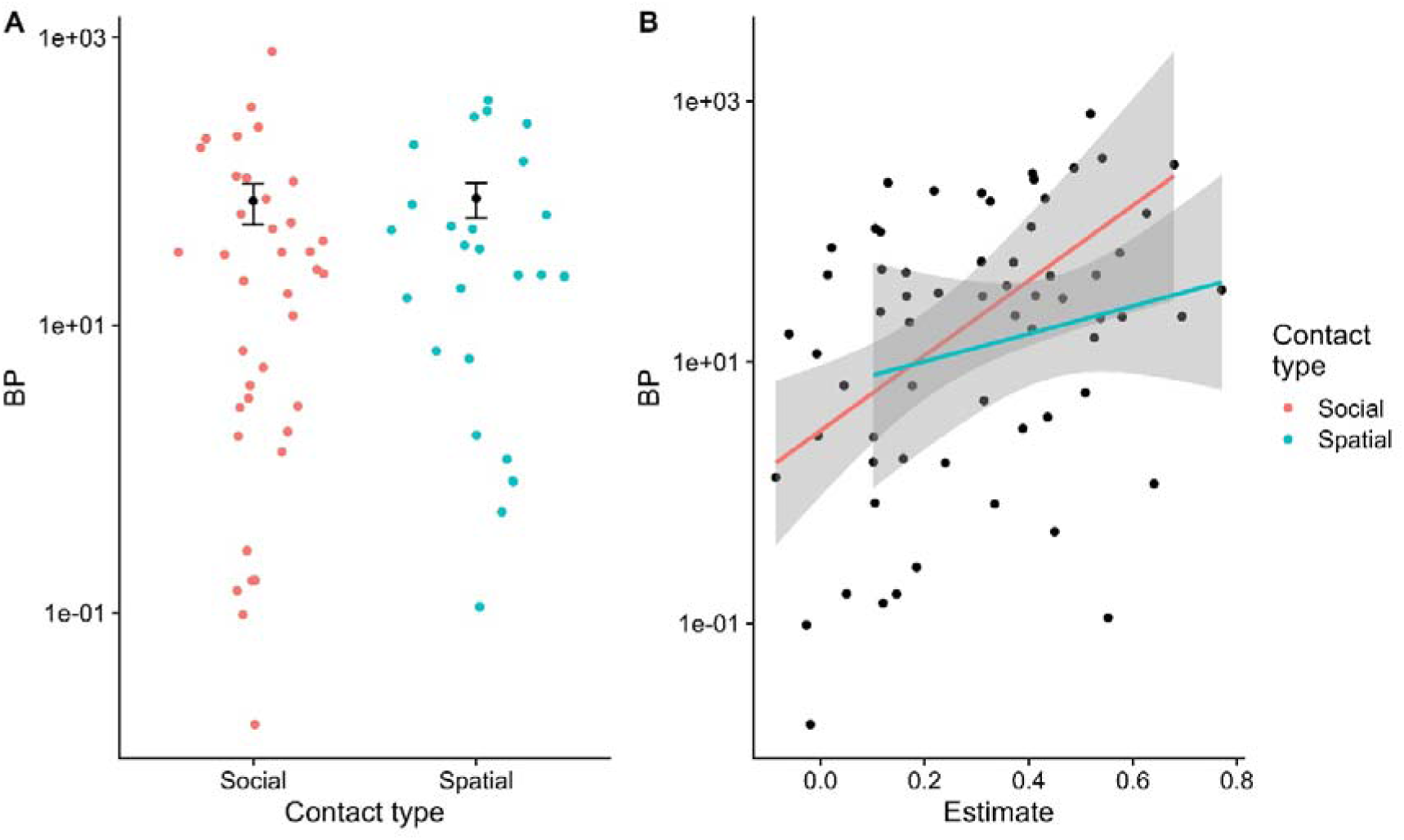
distribution of the Breusch-Pagan Test Statistic as a measure of heteroskedasticity (y axis) according to the type of contact being analysed (panel A) and the magnitude of the linear effect estimate (panel B). The y axis is log-transformed. These findings demonstrate no substantial difference in the levels of heteroskedasticity in the two contact types that might explain spatial models exhibiting a steeper density effect.

**Supplementary Figure 5:**
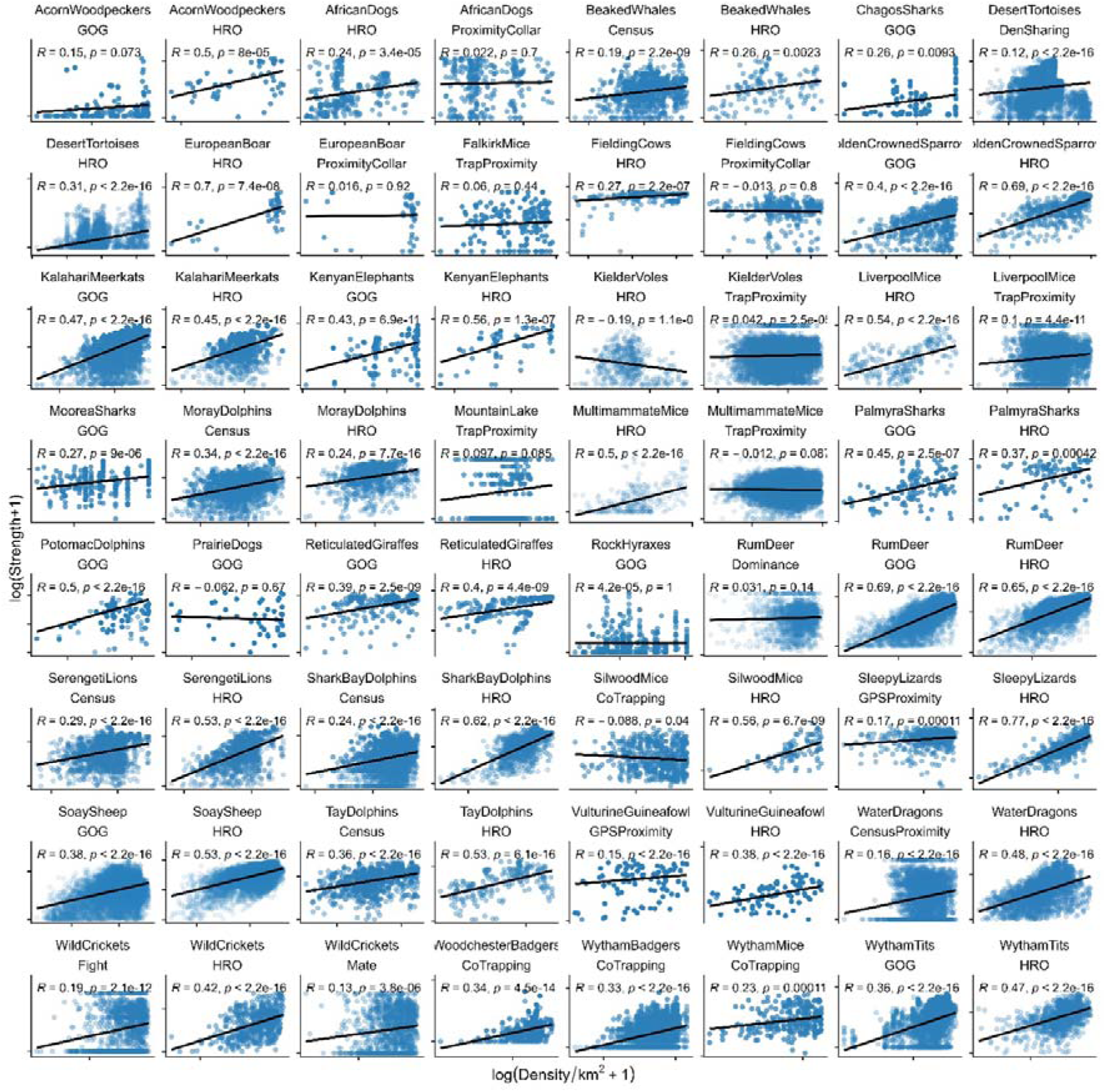
Log-linear relationships between density and network connectedness across animal systems. Density in individuals per area is on the x axis; network connectedness (strength centrality) is on the y axis, both of which are then log(x+1)-transformed. Each point represents an individual-year-behaviour replicate; the lines portray the model fit from our linear models for meta-analysis. Points are semi-transparent to enhance visibility. Panels are arranged phylogenetically following the tree displayed in Figure 2A; GOG=gambit of the group; HRO=home range overlap.

## Supplementary discussion

**Operational implications:** Operationally, the common nature of saturating density effects will impact researchers’ ability to detect density dependence: that is, density dependence could be harder to observe in higher-density areas given the shallower slopes we observed. Most of the systems in this study are relatively long-term studies of known individuals; these populations tend to be in carefully selected, high-density areas that make it convenient to study the focal animal with relatively low operational costs. For example, it has previously been noted that the badgers of Wytham Wood, the red deer of the Isle of Rum, and the Soay sheep of St Kilda are all at high densities for their respective species [37–39]. As such, we may be inherently investigating the upper end of density-connectedness relationships in the wild, and it could be difficult and costly to investigate the effects of low density so widely.

**Limitations:** We acknowledge several limitations of our study, which we nevertheless believe could be remedied in the future. First, many of our social networks were formed of general spatiotemporal associations, and relatively few from specific social interactions – particularly those involving direct physical contact (only 3/36 social networks). Our current dataset could therefore benefit from supplementation with a broader range of direct interactions, particularly involving antagonism or bonding. For example, datasets concerning aggression or dominance interactions (e.g. [40]) or grooming alongside spatial behaviour could inform how density dependence affects the transmission of certain parasites such as mycobacteria [41] or tattoo skin disease [42]. The meta-dataset was also unevenly distributed across animal taxa (Figure 1): there were no primates or bony fishes and only one invertebrate, while rodents and ungulates were over-represented. These biases likely emerge through differences in data collection approaches: for example, although primate social behaviour is often studied with observations of direct interactions that could augment our data as described above (e.g. [43]), the spatial data required to build density distributions are rarely collected in these systems. This is linked to their social structures: our workflow was best suited to studies of fission-fusion societies or relatively asocial animals, rather than those with wide-ranging fixed social groups that are more common in primate systems. Finally, given that our data were observational, we could not account for (or estimate) bidirectional causality between density and social relationships (point 4 in Figure 1): that is, as well as encountering more conspecifics in areas of high density, individuals may be drawn to conspecifics, *creating* areas of high density [44]. To do so might require creating in-depth, high-resolution models of animal movement and group formation (e.g. [45]), potentially making use of telemetry approaches and drawing from large-scale open movement repositories like Movebank [46]. Using remote and next-generation approaches may help to access and incorporate more remote areas, which could also help to ameliorate the substantial geographic biases in our meta-dataset (Figure 2).

**Analytical expansions**: Aside from incorporating more specific interaction types, there exist a range of potential extensions to our analysis. For example, density dependence often varies between age or sex classes (e.g. [47,48]), and age effects on infection are common and ecologically important [49,50], as are sex differences [51]. We chose not to analyse how individual animals’ traits alter the shape or slope of density’s effects for brevity and simplicity; however, given that many of the systems nevertheless include these data, future analyses could make use of this meta-dataset to investigate how density affects connectedness of different classes of hosts. Further, researchers could investigate other behavioural questions such as the role of observation biases; the factors influencing the correlation between spatial and social networks; and the role of environmental drivers and spatial autocorrelation in driving observed patterns of connectivity [44,52]. Finally, as our analysis approximated density-contact relationships and not host-parasite interactions specifically, important future work could investigate whether contact rates (as approximated by network connectedness) ultimately translate to greater infection risk or parasite burden. Although some previous investigations have linked density-related metrics to aspects of infection [53,54], density covaries with a range of other factors including nutrition, cooperation, and competition, all of which could complicate density-driven increases in exposure [55]. For example, in the case of ectoparasite transmission, although contact rates in general would likely increase with density, so too might grooming behaviours that remove parasites; in cases such as these, density’s overall effect on ectoparasite infection may be neutral. In the future, verifying that within- and between-population variation in density-contact relationships translate to variation in infection – and whether these trends might be influenced by flexible avoidance behaviours [56] – will be a vital part of understanding and predicting density-dependent disease dynamics.

## Supplementary acknowledgements

### Acorn woodpeckers

Sahas Barve and Eric L Walters were supported by funding from the National Science Foundation (Grant IOS-1455900). We would like to thank Walt Koenig, Jen Hunter, and many graduate students and field assistants who have helped contribute to this project and the Hastings Natural History Reservation for logistical support.

### African dogs

This system acknowledges support from The Carter Center and the Natural Environment Research Council. Ciro Cattuto and Laura Ozella acknowledge partial support from the Lagrange Project of the ISI Foundation funded by CRT Foundation.

### Bottlenose whales

Fieldwork and data management for this population were supported by Fisheries and Oceans Canada.

### Chagos sharks

This population was funded by the Bertarelli Foundation. Ciro Cattuto and Laura Ozella acknowledge partial support from the Lagrange Project of the ISI Foundation funded by CRT Foundation.

### Cornish cattle

This project acknowledges support from the Biotechnology and Biological Sciences Research Council.

### European boar

Alisa Klamm (AK) is grateful to the Thuringian Ministry of Environmental, Energy and Nature Conservation and the Thuringian Ministry for Infrastructure and Agriculture for funding the research project on wild boar in the Hainich National Park. AK also thanks the people involved in the fieldwork and the copartners: Landesjagdverband Thüringen e.V. and ThüringenForst-Anstalt öffentlichen Rechts. Kevin Morelle (KM) was supported by the H2020 project VACDIVA (grant ID: 862874).

### Golden-crowned sparrows

The research was supported by an NSF Graduate Research Fellowship to Theadora A Block, a CAREER grant (NSF IOS-1750606) and National Geographic grant (WW-R012-17) to Dai Shizuka, and a Laboratoire d’Excellence grant (TULIP, ANR-10-LABX-41) and ANR-SoCo to Alexis S Chaine.

### Firth of Tay and Moray Dolphins

BJC and PSH thank all the colleagues who have helped collect and analyse the Moray Firth and Firth of Tay bottlenose dolphin data. Funding to support these long-term projects was provided by NatureScot, Beatrice Offshore Windfarm Ltd., Moray East and Moray West Offshore Wind Farms, Vattenfall, Seagreen, Inch Cape Offshore Ltd, NnG Offshore Wind, Marine Scotland, The Crown Estate, Highlands and Islands Enterprise, BES, ASAB, Greenpeace Environmental Trust, Scottish Government, Whale and Dolphin Conservation, Talisman Energy (UK) Ltd., Department of Energy and Climate Change, Chevron, Natural Environment Research Council and the Universities of Aberdeen and St Andrews.

### Kalahari meerkats

This analysis uses data on meerkats collected on a project supported by European Research Council Grants numbers 742808 and 294494.

### Kenyan elephants

We would like to thank the Kenyan Office of the President and Kenya Wildlife Service (KWS) for permission to conduct this research and Felix Mdamu and David Korir for help collecting the data. KWS funded radio collars, tracking equipment, and aerial surveys. The research was approved by the Institutional Animal Care and Use Committee (IACUC) of the University of California, Davis, protocol #10087 and received the permission of the Kenya Ministry of Education, Science, and Technology (Permit No. MOEST 13/001/34C 301). Funding for this work was provided by The Lincoln Park Zoo, International Elephant Foundation, and UC Davis Animal Behavior Graduate Group.

### Liverpool wood mice

Data were collected under funding from NERC (grants: NE/G007349/1, NE/G006830/1, NE/I024038/1 and NE/I026367/1), and data collectors: Becci Barber, Trevor Jones, Łukasz Łukomski, Stephen Price, Susan Withenshaw, Niamh Quinn and many undergraduate assistants.

### Mountain Lake mice

Andrea Graham was funded by Defense Advanced Research Projects Agency.

### Potomac dolphins

This population was funded by the Earth Commons Institute, Georgetown University, and under National Geographic Grant WW-022ER-17.

### Prairie dogs

We would like to respectfully acknowledge that our research on prairie dogs occurred on Treaty 4 territory, the traditional territory of the Očhéthi Šakówiŋ, Niitsítpiis-stahkoii ⍰⍰⍰⍰⍰⍰⍰ ⍰⍰⍰⍰, and the homeland of the Michif Piyii (Métis) Nation. We are very thankful to our field technicians for assistance with data collection. We thank S. Liccioli, H. Facette, and the staff at Grasslands National Park for their assistance with permitting and logistical support. This research was supported by a Natural Sciences and Engineering Research Council Discovery Grant RGPIN 04093-2014 (J.E.L.), Parks Canada Contribution Agreement GC-794 (J.E.L.), Saskatchewan Fish and Wildlife Development Fund Student Research Award (J.M.K., and Colleen Crill Matzke), Nature Saskatchewan (J.M.K), and the University of Saskatchewan.

### Rum deer

The authors thank NatureScot for permission to work on the Isle of Rum National Nature Reserve. The authors thank F. Guinness, M. Baker, A. Morris, S. Morris, and many others for collecting field data, and S. Albon and L Kruuk for their contributions to the long-term project. The project has been funded principally by the UK Natural Environment Research Council.

### Serengeti lions

This study population acknowledges NSF grant DEB-1020479.

### Shark Bay dolphins

This study population was funded by NSF grants 2146995, 2106909, 1755229, and 1559380.

### Sleepy lizards

We thank Ron and Leona Clark and Chris Mosey for access to their land and use of the homestead at Bundey Bore Station. We thank Jess Clayton and Dale Burzacott for logistical support, and we are especially grateful to the late Professor Michael Bull and the late Dale Burzacott, who established the sleepy lizard monitoring project. We are also grateful for many students that worked on the sleepy lizard system along the years. (not sure naming them all is helpfull here, so i suggest leaving as is). This work was funded by National Science Foundation of the United States Grant DEB-1456730 and Australian Research Council Grants DP0877384 and DP130100145.

### Soay sheep

The authors thank the National Trust for Scotland for permission to work on St Kilda, and QinetiQ and Kilda Cruises for logistical support. The authors thank J. Pilkington, I. Stevenson and many others for collecting the field data and D. Nussey, J. Slate and M. Morrissey for their contributions to the long-term project. The study on St Kilda has been funded principally by the UK Natural Environment Research Council.

### Vulturine guineafowl

The vulturine guineafowl work was funded the European Research Council (ERC) under the European Union’s Horizon 2020 research and innovation programme (grant agreement number 850859) and Eccellenza Professorship Grant of the Swiss National Science Foundation (grant number PCEFP3_187058) awarded to DRF.

### Wild crickets

WildCrickets would like to thank Luis Rodríguez and María del Carmen Muñoz for unconditional support and providing access to facilities including the WildCrickets study meadow. The following people contributed to video processing and data recording: David Fisher, Ian Skicko, Xing P. Liu, Thor Veen, Carlos Rodríguez del Valle, Alan Rees, Sophie Haugland Pedersen, Hannah Hudson, Jasmine Jenkin, Lauren Morse, Emma Rogan, Emelia Hiorns, Sarah Callow, Jamie Barnes, Chloe Mnatzaganian, Olivia Pearson, Adèle James, Robin Brown, Chris Shipway, Luke Meadows and Peter Efstratiou. The WildCrickets project was supported by the Natural Environment Research Council (NERC); standard grants: NE/E005403/1, NE/H02364X/1, NE/L003635/1, NE/R000328/1, NE/V000772/1 and the Leverhulme Trust.

### Woodchester badgers

This study was funded by the UK Department for Environment, Farming and Rural Affairs, Animal and Plant Health Agency, and the Natural Environmental Research Council.

### Wytham great tits

The Wytham great tit data were funded by ERC AdG 250164, NERC NE/S010335/1 and NE/D011744/1, and collected by the members of the Wytham Social Network Group.

## Supplementary methods – Study System Details

### Acorn woodpeckers

We studied acorn woodpeckers at Hastings Natural History Reservation (36.387° N, 22 121.551° W) in central coastal California, USA. Adults on their natal territory with their social (and genetic) parents were categorized as nonbreeding helpers. Group members living outside their natal territories, or living with birds of the opposite sex that were nonrelatives, were considered putative breeders^1^. Extra-group mating, as well as incestuous mating, is rare in this study population^1,2^. From 1973 to 2021, the majority of the woodpecker population was color-banded (N = 6572 total individuals) and monitored continuously for group size and composition. Each year, territory quality was assigned to each acorn woodpecker social group territory based on the size of the group’s granary (1: <1000 storage holes [low quality], 2: 1000–2500 [medium quality], 3: >2500 [high quality])^3^.

Acorn woodpeckers were caught opportunistically and fitted with dorsally mounted solar-powered nanotags^4^ with leg loop harnesses adjusted for body size^5^. All tags weighed less than 1% of each bird’s body mass. Radio-tagged birds (N=87) were detected by an array of 43 permanently installed autonomous, solar-powered base stations during daylight hours^6^ for a total of 497 days between August 2019 and July 2021.

Tags produced an encoded 64-bit, 2.5 ms radio ping every 1.5 s during the day, even in cloudy weather. Each detection of an individual at the base station was accompanied with a date, time, and signal-strength stamp. All detections were stored in files created every 15 min and stored on removable memory drives^7^.

The raw data were first cleaned to retain only those detections that occurred with high signal strength (within ∼100 m of the receiver). Next, we calculated time spent by each bird (N=87) at each receiver station (N=43). For this, we first partitioned each day into 1-min bins based on the first and the last detection of the day. Next, we assigned a bird to a receiver for a given 1-min bin if we detected that bird more than 10 times at a particular receiver within that minute. This resulted in a cumulative 821,262 minutes of bird presence at receivers.

We classified birds as “associating” if they were detected in the same 1-min bin at the same receiver on a given day. We classified associations as at “home” if the receiver was within 200 m of the bird’s territory or “away” if the bird was detected with high signal strength at a receiver farther than 200 m from its own territory. We detected 175,368 minutes of association between birds (range 2–16 birds associating). Most common associations were between two birds (60,299 minutes of association) and least common association was among 16 birds (only one instance detected). Seventy-eight birds were detected at territories away from their home territory. Distance away from home was highly variable (mean ± SD 483.5 m ± 419 m, range 207–2898 m).

## African dogs

Field site: Dogs (*Canis familiaris*) in rural Chad were collared during the dry (5th March - 17th May 2018) and wet seasons (3rd August – 17th October 2018). Dog owners from six villages participated in the study; Medegue (11°01’48.8” N; 15°26’37.7” E), Ngakedji (9°11’16.5” N; 18°18’10.7” E), Kira (9°10’50.8”N; 18°17’00.3”E), Bembaya (9°11’33.6”N; 18°17’42.3”E), Marabodokouya (9°19’42.3” N; 18°43’20.0” E) and Tarangara; (9°08’19.8” N; 18°42’00.9” E)^1,2^.

Spatial data: Collars were fitted with an iGotU GT600 GPS logger (Mobile Action Technology, New Taipei City, Taiwan). The fix interval was set to 10 minutes. In both the dry and wet field season, an initial two-week deployment was immediately followed by a second, longer deployment using new collars fitted with a modified GT600 unit with a larger battery. Spatial data were cleaned by removing locations taken up to 12 h after the collar was deployed and 12 h before collar recovery. Fixes with speeds greater than 20 km/h were removed and data were discarded for periods when dogs were known to be tied up.

Proximity sensor data: Dog collars were also fitted with a proximity sensor developed by the OpenBeacon project (http://www.openbeacon.org/) and the SocioPatterns collaboration consortium (http://www.sociopatterns.org/). The sensors exchange multiple radio packets per second and proximity is calculated by the attenuation; difference between the received and transmitted power^3–5^. An attenuation threshold of −70dbm was used, detecting encounters within 1–1.5m^5,6^. Inter-logger variability was assessed by comparing the number of packets emitted and received for all pairs of sensors. Data were discarded if there was evidence for deviations from the expected linear relationship between emitted and received radio packets. A contact event was identified when sensors exchanged radio packets for a minimum of 20 consecutive seconds. A contact event ended when the exchange of radio packets ceased in the subsequent 20s period.

## Bottlenose whales

Fieldwork was conducted during most summers between 1988 and 2023, and was nearly always done from a 12m sailing vessel, though other small vessels were used for brief periods of time. The vessel carried out non-systematic surveys of key northern bottlenose whale (*Hyperoodon ampullatus*) habitat on the Scotian Shelf, mostly focusing on the Gully, a large submarine canyon ∼200km from the coast of Nova Scotia. The geographical distribution of effort within the Gully was haphazard, and varied between years. In a subset of years, smaller amounts of time were also spent in the adjacent Shortland and Haldimand canyons. For this analysis we consider observations from the Gully only, as data for adjacent canyons is likely too limited to estimate meaningful variation in spatial behaviour across individuals.

Latitude and longitude for each encounter with northern bottlenose whales, or “sighting”, were measured using various technologies over the course of the long-term study. Loran-C was used from 1988-1992, accurate within ∼400 m^1^, followed by various iterations of increasingly accurate GPS. During sightings, northern bottlenose whales were photographed and photo-identification of individuals was done based on distinctive markings of the dorsal fin. Film photography was used initially, with digital cameras being phased in starting in 2007. IDs are generally side-specific (i.e., left or right), except for those individuals with very distinctive fins that can be unambiguously recognized from either side. For this and most other analyses, we restrict our focus to left-sided IDs only, not considering any right-sided photographs. All observations included in this dataset were from high-quality photographs (quality ratings “3” or “4” out of 4). Additional information can be found in the publicly available photo-identification guide for the northern bottlenose whale project^2^.

Sightings of northern bottlenose whales were considered independent social events. Photo-identifications were linked to the nearest sighting by date and time, from which we drew latitude and longitude data. Any photographs lacking a sighting record within 60 minutes were excluded from further analysis. This primarily included a smaller sets of photographs provided by other research vessels for which we lacked comparable location data. This resulted in 1199 groups over 23 years and a total of 615 unique individuals. Many individuals were seen just once (N = 199). On average, individuals were observed in 5 groups, range 1-66. Observations of groups lasted 10.8 minutes on average (time between first and last photo-identification), with a range of 0-107 minutes.

## Chagos sharks

Grey reef sharks (Carcharhinus amblyrhynchos) have a broad Indo-Pacific distribution and are one of the most common elasmobranch predators on the reefs found across the Chagos Archipelago (6°00′S 71°30′E / 6.000°S 71.500°E). This study utilised acoustic telemetry to track the movements and distribution of sharks across 93 monitoring locations centred around areas of ecological interest (e.g. islands, islets, atolls and seamounts). Information on the monitoring array and tagging procedure can be found in Jacoby et al. 2020^1^.

While tracking of this species occurred continuously between 2013 – 2021, data for this analysis were taken from 2014-2016 when tags at liberty, and thus density of individuals monitored were highest. Tags (Innovasea V16, 69 kHz coded transmitters) acoustically transmit a unique ID code at regular intervals (nominal delay of either 30–90 s or 60–180 s) for the duration of their battery life (∼10 years). Tagged animals were detected whenever they came within range (∼500 m) of an acoustic receiver. Spatio-temporal co-occurrences of tagged sharks were extracted from the telemetry data stream using a Gaussian mixture modelling approach ^2^, and implemented using the ‘gmmevents’ function in the R package asnipe ^3^. Group-by-individual (GBI) matrices that reflected all associations between tagged individuals within a year, were extracted from the model, alongside information on the timing and location of group to the nearest acoustic receiver location. The response variable of social network position and the associated geographic location of social behaviour were derived from these GBI matrices. A total of 142 individuals could contribute to a group throughout the duration of the study.

## Cornish cattle

### Farms and cattle management

Proximity and GPS data were collected from 8 groups of dairy cows on 8 commercial farms in South-West England in Summer and Autumn of 2018 for approximately 7 days per farm. One group consisted of dry cows, whereas all other groups were lactating cows. Some devices malfunctioned and therefore we were not able to obtain data for all cattle in the group (60-91% of the group recorded^1^). Grazing management included rotational grazing, strip grazing, set stocking and free range^1^. Mostly, groups were kept on pasture and brought into buildings only for milking (twice daily), except one group were housed at night, one group were allowed access to buildings at all times and were kept in for two nights and days in the middle of the study period due to inclement weather and one group were allowed free access to all pasture, cubicle housing, and the automated milking system at all times during the study.

### Equipment

Nylon cattle collars with a plastic clasp (Suevia Haiges, Germany) were fitted with a proximity device and a GPS logger such that one device lay at either side of the animal’s neck. The GPS loggers (i-GotU GT-120 and GT-600 devices, Mobile Action Technology Inc., Taiwan) recorded fixes every ten minutes. The proximity device is based on a design by the OpenBeacon project (http://www.openbeacon.org/) and the SocioPatterns collaboration consortium (http://www.sociopatterns.org/) and has been used in contact studies of humans, horses, dogs, and sheep ^2–5^. The proximity sensors exchange low-power radio packets in a peer-to-peer fashion, and this exchange of radio packets is used as a proxy for the proximity of individuals wearing the sensors. To estimate distance between devices, the attenuation of the signals with distance is computed as the difference between the received and transmitted power ^2^. A contact event occurs if at least one data packet is exchanged during a continuous 20-second time window, and a contact is considered broken if no packets are exchanged in a 20-second period ^6^, therefore, contact durations were measured in 20-second blocks.

## Desert tortoises

We monitored tortoise movements using radio telemetry across multiple study sites spanning a 15-year period. Datasets were collected from nine study sites across desert tortoise habitat in the Mojave Desert of California, Nevada, and Utah from 1996 to 2014. Each site was monitored over multiple years, though not all sites were surveyed every year. Individuals were tracked at least weekly during their active season and at least monthly during winter months. Tortoises were fitted with very high frequency (VHF) radio-transmitters (e.g., Model RI-2B [13.8–15.0 g], Holohil Systems Ltd., Carp, Ontario, Canada, or AVM models G3, SB2, or SB2-RL, AVM Colfax CA for older studies), which were attached following established methods (Boarman et al. 1998). Locations were determined using hand-held VHF receivers (e.g., Telonics TR-2, Mesa, AZ, or ICOM RC 10) and recorded with GPS units (Universal Transverse Mercator, North American Datum 1983). Periods of intensive tracking (i.e., multiple relocations per day) were conducted to obtain detailed habitat use data during peak activity for some studies.

Each tortoise was uniquely identified with a numbered paper tag sealed with clear epoxy and permanent notches on the marginal scutes (Cagle 1939). During each encounter, we recorded the individual’s identification number, date and time of observation, GPS location, microhabitat type (e.g., vegetation, pallet, or burrow), and any visible signs of injury or disease. Burrow use was documented by assigning a unique identification number to each burrow, with new IDs assigned when previously unmarked burrows were encountered. Den sharing was used as the contact type, where observations of individuals witnessed within the same den in the same sampling date were taken to be connected.

The dataset was based on monitoring tagged individuals, so data collection was not blind. While most tortoises were monitored consistently throughout the study, logistical constraints and equipment failures occasionally altered the telemetry schedule.

## Ein Gedi hyraxes

The rock hyrax (Procavia capensis) population has been studied since 1999 at the Ein Gedi Nature Reserve (31°28′N, 35°24′E) in Israel (e.g. 1–4). Data used in this study were collected continuously for the years 2000–2020, totaling approximately 75,000 observations. The research focuses on two deep gorges, David and Arugot, located on the western side of the Dead Sea. Each field season, beginning in March and lasting between three and six months, rock hyraxesare trapped and observed daily. Tomahawk live box traps were deployed in secure, shaded locations, baited with cabbage and kohlrabi. Given that rock hyraxes are diurnal, the traps are opened during a fixed morning interval. Once captured, the animals are anesthetized with ketamine hydrochloride (0.1umg/kg) and receive a subcutaneous microchip (DataMars SA) along with either an ear tag (for pups and juveniles) or a lightweight collar (weighing less than 5ug) bearing identification marks. Each individual is then sexed, weighed, and measured, and is given 90 to 150 minutes to recover from the anesthesia. Animals that are recaptured are not anesthetized; they are simply weighed and are promptly released.

Hyrax activity was monitored each day during the field season using 10×42 binoculars and a telescope with 50–100× magnification. Observations were conducted in the early morning, from first light until noon, the time when the hyraxes retreat to their shelters. Every day, a randomly chosen focal group was followed. Although hyraxes primarily spend their time foraging and resting, which allows for the simultaneous monitoring of several individuals, limitations in visibility (due to rocks, trees, and bushes) and the challenge of following up to ten individuals meant that we could not capture the precise duration of every pairwise interaction. Consequently, we recorded interactions on a daily basis: if two individuals were seen interacting at any point during a day, that day was noted as an interaction event regardless of its length. In addition, we recorded the location of each observed hyrax, up to 5m accuracy.

Our annual marking efficiency was high, with about 95% ± 0.5 of group members identified, thus minimizing bias in assessing each group’s social structure. Social interactions that involved unmarked individuals were excluded from analysis. We define positive interactions as those involving direct physical contact (such as huddling or sharing a sleeping burrow) or coordinated behavior (moving together closely or sitting side by side).

## European boar

Behavioral data for wild boar were collected from three European study areas: Bavarian Forest National Park (BFNP, 48°59′N, 13°23′E, Germany), Hainich National Park (HNP, 51°04′N, 10°26′E, Germany), and Kostelec nad Černými lesy (KNC, 50°0′N, 14°50′E, Czech Republic).

BFNP data were collected between October 2021 and January 2023 as part of a research project on movement ecology and African swine fever (ASF) transmission dynamics 1. The study spanned the 250 km² BFNP and the adjacent 684 km² Šumava National Park in the Czech Republic, with elevations ranging from 570 to 1453 m. The landscape consists of mixed coniferous and mountain forests. A total of 42 wild boar were GPS-collared (Vertex Plus, Vectronic Aerospace GmbH, Berlin, Germany) to track movement patterns, home range sizes, and habitat use. Animals were captured using ∼30 m² wood-clad corral traps equipped with live-monitoring cameras and counterweight-triggered gates. The captured individuals were restrained in a net tunnel, handled for approximately 5 min, and released. Ethical approval was granted by the Upper Bavaria government (permit ROB-55.2-2532.Vet-02-20-149).

HNP data were collected between April 2017 and August 2019 to investigate the effects of no-hunting zones on wild boar movement and space use 2. The 25,000 ha study area comprises 54.6% agricultural land, 34.8% forests, 7.3% open land, 3.2% anthropogenic areas, and 0.4% water bodies, with elevations ranging from 180 to 500 m a.s.l. HNP is 75 km² large, with 33 km² designated as a no-hunting zone, including UNESCO-listed primeval beech forests and former military sites where hunting is prohibited. A total of 63 wild boar were GPS-collared (Vertex Lite, Vectronic Aerospace GmbH, Berlin, Germany), with GPS fixes recorded at 30-minute intervals.

KNC data were collected between April 2019 and October 2022 as part of a study on wild boar movement ecology, social behavior, and responses to ASF control measures (including hunting pressure, artificial food supply, and movement barriers–electric and odor fences, and sound traps) 3,4. The 2,900 ha study area, located ∼30 km east of Prague, comprises mixed forests, agricultural land, water bodies, and built-up areas, with an average elevation of 430 m a.s.l. Managed by the Czech University of Life Sciences Prague (Lesy ČZU), this area is heavily frequented for tourism, forestry, and hunting. A total of 84 wild boar were captured using wooden corral traps baited with corn and immobilized via anesthetic darts delivered by airguns. Each individual was GPS/GSM-collared (Vectronic Aerospace GmbH), with fixes recorded every 30–60 min. Ethical approval was granted by the Ministry of the Environment of the Czech Republic (permit MZP/2019/630/361).

For all three datasets, social networks were constructed using the gambit of the group approach, where individuals were considered to be associating if they were within 100 m of each other within a 10-minute interval. GPS data were processed using the spatsoc package 5,6 in R, with group_times used to assign temporal groupings and edge_dist used to calculate pairwise distances between individuals, generating an edge list of proximity-based associations.

## Falkirk wood mice

Trapping for this study took place from 2015-2017 in Callendar Wood (55.990470, −3.766636; Falkirk, Scotland), a 100ha broadleaf woodland, which contain a populations of wood mice, which are naturally exposed to and infected with a wide range of parasites and pathogens. The experiment had three temporal replicates; all of which took place during the wood mouse breeding season: (i) May - July 2015 (ii) June - August 2016 and (iii) July-November 2017. Experimental design for these experiments included randomised nutrition supplementation (high quality food pellets vs unmanipulated) at the grid level and randomised parasite treatment (anthelmintic treatment vs water control) at the individual level within grids. We used a weight-adjusted of 2ml/g dose of both Pyrantel pamoate (Strongid-P, 100 mg/kg) and Ivermectin (Eqvalan, 9.4mg/kg). In 2017, treatment was re-administered 4 weeks after initial dose for all individuals still in the experiment. All animal work was carried out under the approved UK Home Office Project License 70/8543 in accordance with the UK Home Office.

Trapping grids were set up with trapping stations 10m apart, and two traps at each stations, as follows: 2015: 3 grids with 8×8 trapping stations; 2016: 4, 6×5 grids; 2017: 4, 7×5 grids. In each year we live-trapped mice for 3 nights/week using Sherman live traps (H.B. Sherman 2×2.5×6.5 inch folding trap, Tallahassee, FL, USA). Trapping protocols followed (Sweeny, Clerc, et al., 2021). Each tagged individual was followed for a period of 12-16 days (2015-2016) or up to 8 weeks (2017). During this time, we collected morphometric data, blood samples, and faecal samples at every capture. Movement was minimal was minimal between grids.

A total of 261 individuals were captured 882 times across this 3 year study. We used the easting and northing location of the trapping stations where individuals were trapped to estimate density and social network metrics. Edges in the social network between individuals (nodes) were defined as unique mice trapped nearby (within one adjacent trap distance by Euclidean distance) in the same trapping night.

## Golden-crowned sparrow

Golden-crowned sparrow flocks have been monitored at the University of California-Santa Cruz Arboretum since 2009 as part of a long-term research study on their winter social behavior (Shizuka et al. 2014; Madsen et al. 2023). Each fall, first-year migrants were fitted with colored plastic and numbered metal leg bands in unique color combinations. Foraging flocks were surveyed throughout the Arboretum by recording the identities of all banded birds in a flock, which was defined as a group of birds foraging within a 5 m radius. Locations of flocks were recorded using a reference map of 10 m x 10 m grid cells (Shizuka et al. 2014). Observations at seed piles were not considered in this definition of foraging flocks and were excluded from our analysis of flocking relationships. To ensure we were observing winter flocking behaviors rather than interactions between transitory individuals on migration, we limited observations to between November 1 (when most birds had arrived from breeding grounds) and March 1 (when winter flocks begin to break apart and birds begin migration). We further limited our sample to birds that had been sighted 3 or more times throughout this period to remove transient individuals. We then inferred social associations using the ‘gambit of the group’. from co-occurrences in foraging flocks and calculated the simple ratio index (SRI) to represent association strength (see Shizuka et al. 2014 for further details).

## Kalahari meerkats

An individual-based study of meerkats has been running at the Kuruman River Reserve in South Africa since 1993 (26°58uS, 21°49uE, Clutton-Brock and Manser 2016). The study area, which covers approximately 50–60 km2, includes a diverse landscape of dry pans, vegetated sand dunes, and arid bushveld that is typical of a South African savannah, where livestock and game farming form the principal land use. Since the start of the project, approximately 15 groups (range = 6 to 21 groups) and around 200 individuals were followed at any one time (mean ± SD per month = 214.5 ± 59.4, range = 101 to 359). Most individuals were born into the study population and were habituated from birth to allow close behavioural observation. Groups were visited 3–4 times per week in the morning and afternoon throughout the study, with data collected on the composition of groups and on the behaviour, reproductive status, social status, and health status of individuals, so that pregnancies, births, deaths, emigrations and immigrations could be enumerated (summarised in Clutton-Brock and Manser 2016). Most individuals in the population were also weighed at each visit by enticing them onto electronic scales with small amounts of hard-boiled egg or water, and while foraging, GPS locations were taken from the center of the group at 15 min intervals, which we use to estimate home ranges. Though the project began in 1993, all the above data were only collected on multiple groups (> 5) simultaneously from 1998 onwards. Most of our analyses therefore cover the breeding seasons from 1998–2023. The only exception is the GPS data, which were collected in the form needed for our analyses from 2002 onwards. Individuals witnessed within the same group in the same sampling date were taken to be connected.

## Kenyan elephants

### Relevant methods from

The relationship between social behavior and habitat familiarity in African elephants (*Loxodonta africana*). Noa Pinter-Wollman, Lynne A. Isbell and Lynette A. Hart. Proc. R. Soc. B (2009) 276, 1009–1014. doi:10.1098/rspb.2008.1538

### Translocation and sightings

During September 2005, 150 African elephants were translocated from Shimba Hills National Reserve on the coast of Kenya (4.08 S to 4.38 S and 39.58 E to 39.38 E) to Tsavo East National Park (2.08S to 3.78S and 38.18E to 39.38 E), a distance of 160 km. This translocation was part of the Kenya Wildlife Service (KWS) effort to decrease human– elephant conflict in the vicinity of Shimba Hills. Twenty elephant groups comprising adult females, juveniles and calves (average group size 6.8) and 20 independent adult males were moved over the course of 32 days. The release site differs ecologically from the source site and is separated from it by dense human population, providing a unique opportunity for examining the social behaviour of the elephants in a novel environment.

During the translocation, all the elephants were tagged with yellow zip ties on their tails to distinguish them from the local Tsavo elephant population. Unique white numbers painted on the translocated elephants’ backs, natural ear marks and tusk shapes were used for individual identification of the translocated elephants (Moss 1996). Elephants’ ages were estimated based on Moss (1996).

The locations, their time and the identities of the translocated and local Tsavo elephants were recorded in Tsavo East for a year post-translocation using a Geko 201 GPS unit (Garmin Ltd., USA). Road transects were conducted using a vehicle four to five times a week, alternating between four routes of similar length (50–70 km) on the existing roads within Tsavo East National Park. A total of 3371 elephant sightings were recorded, of which 386 and 2985 were the translocated and local elephants, respectively. Of the 150 elephants translocated, data on 83 were obtained, and are presented here. Because males leave the social unit in which they were born at the age of 15, and because the social behaviour of these independent males differs from that of females and their young offspring (Moss & Poole 1983), such translocated males were excluded from our analyses.

### Social association

Elephants were defined as associating with one another if they were sighted within 500 m from one another within a 2 hour time period, based on McComb et al. (2000, 2003). They showed that elephants can individually recognize conspecifics’ vocalizations over great distances (1 km). Therefore, the definition of social association used here includes not only direct interactions but also recognizes the communicative capabilities of elephants to acquire information about the number and identities (translocated or local) of vocalizing conspecifics (McComb et al. 2000, 2003). Thus, the definition of social association used here allows for the acquisition of inadvertent social information about the new environment (Danchin et al. 2004).

## Kielder voles

Briefly, we monitored a natural population of field voles (*Microtus agrestis*) in Kielder Forest, Northumberland, UK across three studies: 2001–2007, 2008–2010 and 2015–2017. Kielder Forest is a man-made spruce forest and field voles are found in grassy clear-cuts. Trapping was performed across different sites, each a forest clear-cut. At each site, Ugglan small mammal traps (Grahnab, Gnosjo, Sweden) were laid out in a grid and checked regularly in ‘primary’ and ‘secondary’ sessions (see below for details). Access to the sites was provided by Forestry England. Newly trapped field voles were injected with a Passive Integrated Transponder (PIT) tag (AVID, UK) for unique identification. At each subsequent capture, we recorded: date, site, trap location and PIT tag ID. This approach allowed us to build up a longitudinal record of the location of tagged voles across multiple sessions. Edges in the social network between individuals (nodes) were defined as unique voles trapped nearby (within one adjacent trap distance by Euclidean distance) in the same trapping night.

Between 2001-2007, trapping was performed from March to November. Primary sessions took place at monthly intervals. At each ‘primary’ session, voles were trapped at 4 different sites. At each site, 100 Ugglan small mammal traps were laid out in a grid spaced 5 m apart. At each ‘primary’ session, traps were checked a total of five times (‘secondary sessions’). More details for the 2001-2007 study are available in^1^.

Between 2008-2010, trapping was performed either from February (2008–2009) or April (2009–2010) to November. Primary sessions took place at monthly intervals. Voles were trapped at a total of 4 different sites – two sites in 2008–2009 and a further two sites in 2009–2010. At each site, 150 Ugglan small mammal traps were laid out in a grid spaced 3–5 m apart. Primary sessions lasted 3 days, and traps were checked a total of five times. More details for the 2008-2010 study are available in^2^.

Between 2015-2017, trapping was performed from March to October. Primary sessions took place at approximately two-week intervals. At each primary session, voles were trapped at 4 different sites. During the study, 3 sites were reassigned due to practical constraints, giving a total of 7 different sampling sites. At each site, 150–197 Ugglan small mammal traps were laid out in a grid spaced 3–5 m apart. Primary sessions lasted 3 days, with traps checked each morning and afternoon. More details for the 2015-2017 study are available in^3^.

## Liverpool wood mice

Live-trapping for this study was carried out in wild wood mouse populations located near Liverpool, UK regularly between May and December for six consecutive years (2009–2014). We sampled 16 trapping grids ranging in size from 2,500 to 10,000 m2, spread across five different woodland sites with 2–3 sites trapped per year. Sites ranged from approximately 2 to 60 km apart. On each grid, trapping stations were placed every 10 m, with two live traps (H.B. Sherman 2 × 2.5 × 6.5 in. folding traps, Tallahassee, FL, USA) at each station baited with grains and bedding material. Full trapping details can be found in (Sweeny, Albery, et al., 2021).

A total of 926 individuals were captured 1,609 times across this 6-year study (max captures per individual = 28, median captures per individual = 4). We used the easting and northing location of the trapping stations where individuals were trapped to estimate density and social network metrics. Edges in the social network between individuals (nodes) were defined as unique mice trapped nearby (within one adjacent trap distance by Euclidean distance) in the same trapping night. Density was calculated using individuals’ centroids in the sampling year based on annual density kernels and using trapping locations.

## Moorea sharks

Owner: Johann Mourier & Serge Planes

Location: Moorea, French Polynesia

The study was conducted at Moorea Island (17°30′S; 149°51′W) in the Society archipelago, French Polynesia. Between 2008 and 2010, a total of seven sites were surveyed on a regular basis along approximately 10 km of the Northern reef of Moorea Island. Among them, six sites were located on the outer reef and were characterized by coral structures from the barrier reef to the drop off (70 m depth) and one site was located inside the lagoon between 2 and 10 m depth within a small channel and characterized by coral patches in a sandy habitat (Mourier et al., 2012; Mourier and Planes 2021). The surveys consisted of 40 min dives (∼30 min dedicated to survey) during which individual blacktip reef sharks were identified by photo-identification, using unique, lifelong color-shape of the dorsal fin (Mourier et al., 2012).

Associations between individuals were defined using the “Gambit of the Group”, assuming that all individuals observed together are then considered as “associated.” An experienced diver conducted a stationary visual census at each site monitored and identifying sharks within a ∼100-m radius area (made possible by the high visibility conditions in these tropical waters). As most sharks usually remained together during the time of the dive, we considered the largest number of individuals observed within a 10-min period to be part of a group. We are confident that observed associations represented true grouping structure because groups were spatio-temporally well-defined and sharks were engaged in specific social behavior (e.g., following, parallel swimming or milling; Mourier et al., 2012). To avoid the potential for weak and nonrelevant associations between pairs of individuals with very low number of sightings, we used a restrictive observation threshold to include only individuals observed more than the median number of sightings (median = 14). Thus, all individuals seen less than 15 times were removed from the analyses to ensure that associations were estimated with high accuracy and precision.

Our dataset totaled 180 groups among 105 adult blacktip reef sharks.

## Moray and Tay dolphins

Data are from a long-term individual-based study of bottlenose dolphins on the east coast of Scotland^1^. Boat-based photo-identification (photo-ID) data have been collected since 1989, initially focussing on an area that was subsequently designated as the Moray Firth Special Area of Conservation (92/43/EEC), with occasional surveys further afield^2,3^. During this time the population expanded its distributional range^4,5^ and since 2009 photo-ID surveys have also been regularly carried out around Tayside and adjacent waters^5,6^. Photo-ID surveys take place annually from May to September, with occasional surveys at other times of the year. Survey routes are chosen to maximise sighting probability while providing wide coverage of the study areas and all surveys were carried out under NatureScot Animal Scientific Licences. Surveys adhere to standardised protocols^2,5^ where photographs were taken of the dorsal fins of as many individual dolphins as possible, graded for quality and using unique markings matched to the Universities of Aberdeen and St Andrews catalogue of known bottlenose dolphins on the Scottish east coast. All dolphins within 100m and engaged in similar activities or travelling in the same direction were considered to be in the same group and associated. Sex was determined using genital photographs or if adults were seen in repeat association with a known calf^7^. Year of birth was estimated from photographs of calves (up to 2 years old) based on their foetal folds, size, colour and behaviour^8^.

Sightings data from this population were provided for this study for every bottlenose dolphin group encountered from 1990 to 2021 including the date, location, estimated group size, unique identifier of each dolphin identified, and their sex and year of birth if known. This dataset comprised sightings of 903 individual bottlenose dolphins (males, females, unknown sex and all ages) in 3071 groups and was split into two study areas, Moray (730 dolphins in 2525 groups) and Tay (339 dolphins in 546 groups). Individuals from this population are known to travel across the east coast of Scotland and 166 dolphins were photographed in both areas.

## Mountain Lake mice

Data for the Mountain Lake Mice were collected as part of two 3-year deworming experiments conducted at the Mountain Lake Biological Station, Pembrooke, VA from 2016-2018 (37°22’N, 80°31W). Eight 0.5 ha grids (8 x 8 traps with 10 m spacing) were live-trapped for 2-3 night sessions approximately every 2 weeks using Sherman live traps (H.B. Sherman 2×2.5×6.5 inch folding trap, Tallahassee, FL, USA) from May to August/early September each year. Trapping protocols followed (Sweeny et al. 2020) with each individual receiving a numbered ear tag upon first capture. Movement among grids was minimal with less than 2% of individuals captured on multiple grids. Thus, grids were treated as separate populations for local density and social network calculations. Edges in the social network between individuals (nodes) were defined as unique mice trapped nearby (within one adjacent trap distance by Euclidean distance) in the same trapping night. Animal care and use protocols were approved by Princeton University, and the research was supported by a DARPA grant to ALG (68255-LS-DRP).

## Multimammate mice

*Mastomys natalensis*, the multimammate mouse, occurs throughout sub-saharan Africa and is an important agricultural pest species as well as host to a number of diseases of human health importance (e.g. plague, Lassa virus and leptospira (Fichet-calvet and Becker-ziaja, 2014; Holt et al., 2006; Mccauley et al., 2015). In Morogoro, Tanzania, where field data for this study was collected, *M. natalensis* undergoes considerable population fluctuations, with densities ranging from 10ha-1 during the breeding season up to 150ha-1 outside the breeding season. Population dynamics have been monitored using a robust design open capture mark recapture on a monthly basis since 1994 and is ongoing (Leirs et al., 2023). For the purposes of this study, data was used for the period of 1994 - 2022. The study design is as follows: 300 single capture Sherman traps were placed on a 300 x 100 m grid and baited with a mixture of peanut butter and cornflour. Trapping occurred for three successive nights; trapped individuals were weighed, sex and sexual condition recorded and individually marked with a unique toe clipping code if it is the first time they were captured. More details about the study site and capture methods can be found in Leirs et. al 2023. Edges in the social network between individuals (nodes) were defined as unique mice trapped nearby (within one adjacent trap distance by Euclidean distance) in the same trapping night.

## Palmyra sharks

We tagged 41 grey reef sharks (*Carcharhinus amblyrhynchos*) that were caught on the forereefs of Palmyra Atoll (5°540 N 162°050 W) in the Central Pacific Ocean. Palmyra is a US Wildlife Refuge and includes large numbers of sharks and other predators (e.g. Bradley et al. 2017). Sharks were caught on hook and line and surgically implanted with Vemco V16 acoustic transmitters (69kHz, semi-randomized delay 60-180 seconds). Transmitters had battery lives of four years and were detected by a network of 55 acoustic receivers (Vemco, VR2W) positioned around the forereef and within the lagoons and back reefs (see Papastamatiou et al. 2018 for full details). Receivers would record the date and time of sharks that swam within range of the receiver (approximately 200-300 m).

We used a gambit of the group approach to generate dynamic social networks, where clusters of sharks co-occurring at receivers through time were identified using Gaussian mixture models (Psorakis et al. 2012, Jacoby et al. 2016). These clusters consisted of individual sharks that visited the same receivers at the same time. Adjacency matrices were produced based on how often dyadic pairs co-occurred within identified clusters. Full model details can be found in Papastamatiou et al. 2020.

## Potomac dolphins

### Field Site and Dolphin Population

The Potomac-Chesapeake Dolphin Project has been conducting annual surveys of wild Tamanend’s bottlenose dolphins (*Tursiops erebennus*) in the Potomac River and mid-Chesapeake Bay since July 2015. Approximately 550 km2 in the lower Potomac and mid-Chesapeake Bay are surveyed annually with 90% of survey effort concentrated in a 58 km2 area. Since the project started, over 2000 individuals have been identified with 36% sighted more than one day and 20% returning to the area more than one year. Depending on one’s definition, the populations using the area are migratory and inhabit Chesapeake mainstem waters and tributaries in the warm months (April-October) before moving south during the cold months..

### Data Collection

All data for this study were collected between 0700 and 1800 between April to October. For the current analyses, survey data annually from 2015 to 2018 were included for a total of 102 surveys with ID data on 1151 individuals (410 were re-sighted). Group sizes ranged from 1-163. The predominant group activity was determined the same way as in Shark Bay. However, group definitions differ slightly due to differences between the two systems. A survey group is defined as a close association or as an aggregation depending on the circumstances. A close association is a discrete group usually comprised of a small number of animals where the 10m chain rule can be used to determine association (as is the case for Shark Bay). An aggregation is a large number of animals that are within a 100m radius and may or may not be delineated into discrete groups as defined by the 10m chain rule. In aggregations, animals are clearly connected to each other with members moving between discrete groups or, in the absence of discrete groups, among each other. Under either circumstance (with or without discrete groups), animals move among each other in such a way that at some point during the survey they have likely been connected by the 10m chain rule.

## Prairie dogs

We studied black-tailed prairied dogs on one colony in Grasslands National Park, Saskatchewan, Canada (49.06N, 107.36W) as a long-term project from 2015 to 2020^1^. Our goal was to capture all individuals to get a full colony census of the age and sex structure within the colony as well as population dynamics. We used Tomahawk traps (Tomahawk, Hazelhurst WI) to capture each individual, and subsequently tagged each pinnae with alphanumeric ear tags (Monel #1, National Band and Tag Co., Newport, Kentucky) and painted their dorsal pelage with a unique symbol for future identification. We weighed, sexed, and aged each prairie dog at first capture.

We conducted our behavioural observations during April (pre-juvenile) and June (post-juvenile emergence) 2017 within a 3-hectare area located centrally within the colony as all animals in this area were marked and readily identifiable from a distance with binoculars. We marked the 3 hectares with a 15 m Cartesian grid system to record the location of individuals during focal observations. As prairie dogs are a sedentary species, space use and social structure are not independent of each other and we recorded both social and movement behaviours. 57 individuals aged 1 year and older were observed across 9 coteries (mean = 6.61 individuals per coterie, range = 2 to 14). We recorded behaviours as we sat > 40 m from marked individuals during peak activity hours for prairie dogs based on season. We recorded all affiliative (sniffing, jump-yipping^2^, greet-kissing, mutual vigilance, shared foraging) and agonistic (fighting, chasing, standoffs, territorial defense) interactions between all individuals in 3-4 hour sessions. We could record all social encounters across all animals as encounter rates were relatively low. We also conducted focal scans to record all activities and locations (<1 m) of each individual to establish their home range. We conducted four 20-min focal scans per individual: two in April and two in June. We recorded an average of 56 locations (range 41-76) for each home range per season. We recorded 380 person-hours of behavioural observation and 872 interactions between known individuals. We created two networks from weighted undirected matrices of behavioural data and home ranges from the focal scan movement data.

## Reticulated giraffes

This study was conducted in Ol Pejeta Conservancy (OPC), a 364 km2 semi-arid wildlife reserve located on the equator (0° N, 36°56’ E) approximately 220 km north of Nairobi, Kenya. All giraffe at OPC were recognized using individually unique spot patterns along their necks, and classified to age groups as described in (1). At the time of this study, OPC had a population of 212 reticulated giraffe. At the conclusion of the study period, OPC’s giraffe population consisted of 160 adults, 20 subadults, 21 juveniles, and 11 neonates. The population exhibited a 50:50 sex ratio.

From January 21 to August 2, 2011, giraffe group composition and membership were recorded for all giraffe groups sighted while driving pre-determined survey routes. Observed giraffe groups were followed off-road until a complete census of the individuals present was accomplished. A group was defined as a set of individuals engaged in the same behavior, or moving in the same direction or toward a common destination, as long as each giraffe was no more than 500 m from at least one other group member. During the study period, a total of 1089 observations of giraffe groups were collected. On average, 30.7 giraffe were observed per day, distributed between four to six groups. Each individual giraffe was observed on average 31.1 times (approximately once per week). A social network was constructed from observed association patterns, using the “gambit of the group” approach.

## Rum deer

Adapted from: Albery GF, Morris A, Morris S, Pemberton JM, Clutton-Brock TH, Nussey DH, Firth JA (2021): Multiple spatial behaviours govern social network positions in a wild ungulate. Ecology Letters 24 (4), 676-686. https://doi.org/10.1111/ele.13684

The study was carried out on an unpredated long-term study population of red deer on the Isle of Rum, Scotland (57°N,6°20′W). The natural history of this matrilineal mammalian system has been studied extensively (Clutton-Brock et al. 1982), and we focussed on females aged 3+ years, as these individuals have the most complete associated census data, and few males live in the study area except during the mating period. Individuals are monitored from birth, providing substantial life history and behavioural data, and >90% of calves are caught and tagged, with tissue samples taken^1^.

Census data were collected for the years 1974-2017, totalling 423,070 census observations. Deer were censused by field workers five times a month, for eight months of the year, along one of two alternating routes^1^. Individuals’ identities, locations (to the nearest 100M), and group membership were recorded. Grouping events were estimated by seasoned field workers according to a variant of the “chain rule” (Castles et al. 2014), where individuals grazing in a contiguous group within close proximity of each other (each individual under ∼10 metres of at least one other individual in the group) were deemed to be associating, with mean 130.4 groups observed per individual across their lifetime (range 6-943). The mortality period falls between Jan-March, when there is least available food, and minimal mortality occurs outside this period. We only used census records in each May-December period, from which we derived annual social network position measures as response variables. We elected to investigate this seasonal period because it stretches from the spring calving period until the beginning of the mortality period, simplifying network construction and avoiding complications arising from mortality events. Our dataset totalled 3356 annual observations among 532 grown females (Figure 1).

We constructed a series of 43 annual social networks using “gambit of the group,” where individuals in the same grouping event (as described above) were taken to be associating^3^, and using dominance interactions.

## Serengeti lions

The research data were collected on the well-studied population of African lions inhabiting a 2700km2 study area within the Serengeti National Park, Tanzania. Multiple rivers and tributaries transect this area, which encompasses both a grassland plains habitat in the southeast, and woodland habitat to the north. There are two main seasons, the wet season spanning November to May, and the dry season spanning June to October. While the woodlands habitat maintains relative stability year-round, the grasslands habitat experiences greater fluctuations in rainfall (Mosser and Packer, 2009). This leads to large seasonal migrations of prey species across the plains, with lower populations densities in the dry season. As such, lion territories shift with the seasons across the landscape, tracking prey species migrations (Packer et al., 2005).

Lionesses exist in egalitarian fission-fusion “prides” composed of related adults and their offspring. In the Serengeti study area these prides can range in size from 2-20 individuals. Within these prides, individuals spend large proportions of time in smaller subgroups, and frequently spend time alone. The composition and size of these subgroups is highly dynamic, fluctuating daily, but female pride-mates usually stay within 5-6 km of each other (within vocal communication range). At maturity (2-3 years of age), approximately 75% of female offspring are recruited into their natal pride, while 25% disperse to form new prides, usually in adjacent territories (Packer, 2023; Pusey and Packer, 1987).

At maturity, all male offspring disperse from their mother’s pride in related groups of brothers and cousins. During this nomadic life stage multiple kin cohorts may form alliances, resulting in “coalitions” of up to nine males. Once they reach full body size (∼ age 4 years old), these coalitions “take over” residence of a pride of females by outcompeting the existing resident male coalition (Packer, 2023; Pusey and Packer, 1987). Male coalition membership does not change once they are resident within a pride. No evidence of a dominance hierarchy has been recorded between male coalition mates (Packer et al., 2001; Packer, 2023).

The data for this study were collected over a 30-years period from 1984 to 2013. One female per pride was fitted with a GPS collar. Each collared lion was located at least once every two weeks, and individuals within 200m of each other were taken as co-occurring. These co-occurrences were recorded as part of one unique sighting event, with GPS coordinates, date and time stamps. Additionally, opportunistic sightings of individuals within the study area were recorded in the same manner. Identification of individuals with a high level of accuracy was possible using facial markings and whisker spots. For each individual lion within a sighting event, age (estimated from the first sighting as a cub or adult), sex, natal pride (if known), and current pride information was also recorded. We constructed social networks based on whether individuals were observed in the same group on the same date.

## Shark Bay Dolphin Research Project

Field Site and Dolphin Population: The Shark Bay Dolphin Research Project has been conducting a longitudinal study of bottlenose dolphins (*Tursiops aduncus*) since 1984, monitoring over 1,900 individuals in the eastern gulf of Shark Bay, Australia. The relatively pristine study area extends 600 km2 east of Peron Peninsula and consists of embayment plains (5–13 m), shallow sand flats and seagrass beds (0.5–5 m), and deeper channels (6– 13 m). Using boat-based sampling, field observers identify individual dolphins using dorsal fin markings and shape, and also matched photographs to a digital identification catalog. Dolphin ages were estimated based on year of birth, the birth of their first calf (Mann et al. 2000; McEntee et al. 2023), or degree of ventral speckling (Krzyszczyk and Mann 2012). Individuals were considered to be adults if they were 12 or more years of age, or for females, if they had given birth to a calf (Mann et al. 2000). Finally, sex was determined by visual observation of the genital area, genetic analysis, or the presence of a dependent calf. Both sexes remain in their natal area (bisexual philopatry) in this residential population.

Data Collection: All data for this study were collected between 0600 and 1900 during all seasons using survey methods.

A survey is a “snapshot” of a group or individual. When dolphins were sighted, researchers instantaneously estimated initial activity and distance from the research vessel before approaching to within 100 m. Once the research vessel was close enough for observers to identify individuals a survey was initiated on all individuals in the group based on a 10 m “chain rule” (Smolker et al. 1992). For each survey, observers performed a scan of all individuals to assess their behavioral state as one of six categories: foraging, resting, socializing, traveling, other, and unknown. A “predominant group activity” was assigned for the first five minutes of the survey based on the activities of at least 50% of the individuals (Mann 1999) for at least 50% of the time. The final dataset included 7293 group sightings of 910 individuals between 2008 and 2019.

## Silwood wood mice

The field data collection was led by Sarah Knowles, and the data was cleaned and processed by Bryony Allen, Sarah Knowles, and Aura Raulo.

Data were collected over a one-year period (Nov 2014–Dec 2015) from a wild population of rodents in a 2.47 ha mixed woodland plot (Nash’s Copse) at Imperial College’s Silwood Park campus, UK. Data for this study was collected as part of a longer-term rodent capture-mark-recapture study, where several rodent species were caught (*Apodemus sylvaticus*, *Apodemus sylvaticus*, *Apodemus flavicollis* and *Myodes glareolus*). Trapping was performed every 2-4 weeks, using 122 small folding Sherman traps (5.1 x 6.4 x 16.5cm, H. B Sherman). Traps baited with eight peanuts, a slice of apple and sterile cotton wool for bedding were set at dusk and collected at dawn, with all animals processed, sampled and then released inside the 100m2 grid cell they were captured in. As part of processing, captured individuals were identified to species, sexed, weighed, and aged (juvenile or adult) based on size and pelage characteristics. At first capture, all individuals were injected subcutaneously with a passive integrated transponder tag (PIT-tag) for permanent identification. Ear punches were collected from all mice and stored in ethanol at −20°C to provide genetic material for host genotyping. Live traps were set in an alternating checkerboard design, to ensure even coverage. Edges in the social network between individuals (nodes) were defined as unique mice trapped nearby (within one adjacent trap distance by Euclidean distance) in the same trapping night.

## Sleepy lizards

The following description is adapted from Payne et al. (2022).

We studied sleepy lizards (*Tiliqua rugosa*), a species of skink native to southern Australia. Sleepy lizards are large (adults are 400-950 g, with snout-vent length 25-35 cm), mainly herbivorous, and can live up to 50 years (Bull, 1995; Bull et al., 2017). Social network studies have shown that sleepy lizards are pair-living with strong male-female pair bonds (Leu et al., 2010, 2011), exhibiting a long-term socially monogamous mating system (Bull et al., 1998; Leu et al., 2015). Sleepy lizards are primarily active during the austral spring, September to December (Bull, 1987). Overnight and during periods of heat stress, they shelter in shaded refugia (i.e., large shrubs, logs, or burrows) (Kerr et al., 2003).

Our field site was an ∼ 1.2 km2 area near Bundey Bore Station (33.888240° S, 139.310718° E), South Australia (about 150 km north of Adelaide). The region has a semi-arid Mediterranean climate. The local site is dominated by chenopod shrubs (e.g., Maireana and Atriplex spp.) and patches of black oak trees (Casuarina spp.), with various annual plants growing between and under these shrubs and trees (e.g., the nonnative Ward’s weed, Carrichtera annua, a preferred food item for sleepy lizards). In most years, food is more abundant in early spring when conditions are relatively cool and wet, becoming much less abundant later when conditions are hotter and drier.

As part of a long-term monitoring study, we used GPS units (horizontal precision +/− 6 m (Leu et al. 2010a)) to record adult lizard movement during their active season (i.e., the austral spring) from 2008 through 2017, excluding 2012. In 2008 through 2014, GPS units (Leu et al., 2010) took one GPS fix per 10 minutes, while in 2015 through 2017, GPS units took one fix per two minutes. To reduce autocorrelation in the GPS data and ensure consistency across years, following (Michelangeli et al., 2021), we thinned the GPS data from all years to one point per 20 minutes. We removed GPS errors according to fix accuracy (e.g., horizontal dilution greater than three), using an algorithm modified from Bjorneraas (2010) – which identifies errors via displacement, speed, and turning angle – and with manual inspection of GPS tracks for obvious errors.

## Soay sheep

This study was carried out on a long-term study population of Soay sheep (*Ovis aries*) on the St Kilda archipelago. The Soay sheep have lived wild on the archipelago for millennia, and have been monitored since 1985 in our study area in Village Bay on Hirta. The natural history and population dynamics of these sheep have been extensively documented (Clutton-Brock & Pemberton, 2004). Sheep are captured, marked at birth, and followed longitudinally across their lifespan to collect morphological, behavioural, parasitological, and immunological data.

Census data were collected for the years 1988-2018. Sheep are censused 30 times per year (10 each in spring, summer, and autumn). Experienced fieldworkers follow established routes noting the identity, spatial location (to nearest 100m OS grid square), behaviour and group membership of individual sheep. Fieldworkers assign individual sheep to groups using a ‘chain rule’, where individuals in close proximity are classed as associating (Castles et al., 2014). In total, census data comprises 357,283 total observations of 81,769 groups. Individuals were associated with a mean of 50 unique groups across our total dataset (range 1-464). We used census records to construct annual social networks for each individual sheep, based on group membership in the same observation event.

## Vulturine guineafowl

The vulturine guineafowl (*Acryllium vulturinum*) is a terrestrial bird species (body size: 60-72 cm) that lives in the arid and semi-arid savanna of East Africa. Vulturine guineafowl are gregarious and typically live in stable groups of 15-65 individuals ^1^. Groups are highly cohesive throughout and across the day. Previous work has suggested that groups are non-territorial, establishing highly overlapped home ranges among neighbouring groups ^2,3^. Groups frequently roost communally ^3^ and when the ecological conditions become harsh (dry seasons and droughts), groups can expand their home range areas ^2,4^ and stable groups merge with preferred groups to form supergroups ^1^.

The vulturine guineafowl project was established in 2016 and collects data on up to 23 social groups (a total of 1189 birds) around the Mpala Research Centre in central Kenya. These data include fitting birds from each group with a global positioning system (GPS; 15g eObs solar) tag. These tags collect 1 data point per second (1 Hz), 10 consecutive data points every 5 minutes, or 1 data point every 15 minutes at the least, depending on the battery conditions (see ^5^ for more details). Here we used data from every 15th minute. In addition to the GPS tags, almost all individuals in our population have been fitted with a unique combination of colour bands for individual identification. Each day (morning and evening), a field team census groups in the area, recording the membership of each observed group and its location.

To obtain group size, we used the data from the group compositions for each month (see ^6^ for details) and combined the GPS data (where groups are) with group membership. In brief, we construct networks of robust co-observations among individuals ^7^, and then apply a community detection algorithm (walktrap community algorithm, using igraph package ^8^ in R) to identify groups. From these groups, we could quantify what GPS tags were in the same group and how many birds the group contained, thereby allowing us to estimate local densities.

The contact network was constructed based on the observation that meeting groups tend to mingle completely; each identified group-group contact event assumed that all individuals in one group were meeting all individuals in another group, with a threshold of 20 metres taken to represent a contact event.

## Water dragons

Data were collected as part of an ongoing behavioural study of a population of eastern water dragons (*Intellagama lesueurii*) inhabiting an urban city park, Roma Street Parklands, Brisbane, Australia (27°27′46′ S, 153°1′11′ E). Consisting of approximately 336 individuals, the population inhabits a highly heterogenous, man-made, curated public garden (Strickland et al, 2014). Behavioural surveys were conducted twice daily (morning 7.30–10.30 and afternoon 13.00–16.00) approximately three days a week from November 2010 onwards. Data collection was restricted from September one year until April the following year, representing the season in which dragons are active and not in brumation.

For each individual encountered, a photo of their left and/or right facial profile, along with their GPS location was recorded. An individual’s sex was assigned based on sexual dimorphism and dichromatism (Thompson, 1993). The individual’s immediate behaviour at the time of the observation was recorded, which included a spectrum from resting to agonism (e.g. head bobbing, tail slapping, arm waving, chasing and physical combat).

Profile photographs taken during surveys were used to identify individuals using a previously established method for this population (Gardiner et al, 2014). This method employed interactive identification software (I3S Spot, v. 4.0.2) which compared individual facial profile scale patterns from images taken during behavioural surveys to an existing photo library. We also took GPS locations of every observation; on each census, individuals with less than 1.85m between them were taken to be in contact.

## Wild crickets

The aim of the project was to monitor a wild population of field crickets (*Gryllus campestris*) living in a meadow in North Spain. This species is annual, and every individual spends most of its life inside or in the vicinity of burrows excavated in the ground. We monitored the natural population for 12 consecutive years. Every year, we found all the burrows in the meadow and marked them with a flag having a unique number. We trapped every individual in the population shortly after they emerged as adults and marked them with a plastic tag with a unique code for each individual. We used up to 140 infra-red video cameras to record 24hrs a day at burrows and an area of about 20cm in diameter around the burrow from the date of first adult emergence to the date when the last adult in the population died. We then watched the video and extracted a number of relevant behaviours for each recorded individual, which were then used to form the social networks. A more detailed description of our methodology can be found in Rodríguez-Muñoz et al. (2019).

## Woodchester badgers

The Woodchester Park badger study is conducted at Woodchester Park, Gloucestershire, UK (51.7°N,2.25°W), and has been ongoing since 1976 ^1,2^. For the purposes of this study co-trapping data were used to estimate both population density and social networks (note that co-trapping will only provide a proxy for social associations in this system;^3^). The study site is divided into three zones of approximately equal size, each of which is trapped four times per year from May to January inclusive (no captures occur in the intervening period to avoid catching dependent cubs and their mothers). Box traps constructed of steel mesh are set close to each active main and outlying sett (detected by sett activity surveys in the build-up to trapping) and baited with peanuts for up to 8udays. Traps are then set for two consecutive nights and checked the following mornings. At each active sett, more traps are deployed than expected to be required. On first capture badgers are permanently marked with a unique ID tattoo on the abdomen allowing them to be identified on future captures ^4^. Individuals are typically caught (on average) approximately two times per year^2^.

Co-trapping networks can then be constructed based on individuals being caught at the same sett on the same day. Because these networks are constructed using co-trapping data only they can include all individuals regardless of their age or sex.

## Wytham badgers

The Wytham Badger Project ran 1987 to 2019 and attempted to monitor all badgers resident in the Wytham Woods SSSI, a 4.24 km2 mixed woodland in southern England (51° 46′ N, 1° 20′ W; for further information, see Macdonald & Newman 2022). The population is situated on a hill surrounded by the River Thames on three sides and the A34 motorway on the fourth, mimimizing migration (immigration/emigration rate = 3%: Macdonald and Newman 2002). Over the study period, every active communal burrow system, termed a “sett”, was trapped using string-trigger traps for two or (most commonly) three nights, three– four times per year, at regular seasonal intervals. Badgers were transferred to holding cages between 7:00 and 9:00 a.m., transported to a central feldstation, and sedated with 0.2 mL ketamine hydrochloride/kg body weight by intramuscular injection (McLaren et al. 2005). On first capture (typically as a cub or yearling), each badger received a unique numerical inguinal tattoo. The population divided into 23 social groups, established from frequent baitmarking surveys (Buesching et al. 2016), where each social group utilized several setts consisting of 1–10 holes, with on average 5.5 individuals cohabitating in the average sett (range 1 to 28).

In total, the study amassed 11,488 captures of 1823 individuals, to which an enhanced Minimum Number Alive (eMNA; Bright Ross et al. 2002) enumeration procedure was applied. Population size averaged 242 adults ± 15.14 SD, range = 222–263) plus 66 cubs (± 8.1 SD, range = 47–97) from 2005 to 2009. Thereafter, following high, and unexplained, mortality across age classes in 2010, the population settled to a slightly lower but stable phase through to 2015, comprising 195 adults (± 17.06 SD, range = 177–217) and 49 cubs (± 15.47 SD, range = 24–66; Bright Ross et al. 2020). In 2019, population density was 44.55 ± 5.37 badgers km-2, where Wytham Woods has consistently had the highest density of European badgers ever recorded (Macdonald and Newman 2022). Co-trapping networks were then constructed based on individuals being caught at the same sett on the same day.

## Wytham tits

The study was conducted in Wytham Woods, Oxford, UK (51°46′ N, 1°20′ W), on a great tit (Parus major) population monitored using standardized protocols since the 1960s [1]. Great tits breed almost exclusively in nest boxes fixed at 1020 GPS-mapped locations across Wytham Woods [2]. Over 98% of breeding great tits occupy a single nest box per year, with breeding spanning April–July and comprising nest building, egg laying, incubation, and offspring rearing. Nestboxes were regularly checked to monitor breeding attempts, identify (or mark) adults (days 6–14 of the nestling phase), and mark nestlings (day 15) with unique British Trust for Ornithology (BTO) metal leg rings, alongside taking standard morphometric measurements [1–3].

In winter (September–March), great tits form highly dynamic, fission-fusion feeding flocks with frequent turnover [3–5]. Since 2007, great tits captured during breeding or winter mist-netting have been fitted with plastic leg rings containing a unique RFID microchip, resulting in the RFID tagging of ∼90% of the population [4]. RFID tags enable tracking of individuals at sunflower feeding stations equipped with two RFID antennae (Dorset ID, Aalten, Netherlands) at placed in 65 stratified-grid locations through in winters starting 2011, 2012, and 2013. During these winter seasons, these feeders were open every weekend from End-Nov to End-February (13 weekends), continuously scanning for RFID-tagged birds from pre-dawn to post-dusk.

### Winter Population Social Structure

RFID detections generated a spatiotemporal datastream reflecting the bursts of activity as flocks arrived and fed. A machine-learning algorithm (Gaussian mixture model) assigned detections to the most likely flock or ‘gathering event’, providing a robust and effective method for determining flock co-membership [5]. From the resulting group-by-individual matrices, social networks (association matrices) were constructed using the Simple Ratio Index [5,6].

For this analysis, non-directional weighted networks were built from the yearly (i.e. whole winter season GBI matrix) to measure dyadic association propensity, and the extensive sampling here reduces the typical limitations of the ‘gambit of the group’ approach [7]. Networks included all RFID-detected individuals linked to a unique BTO ring number. Due to high turnover and movement, kin structure was weak, with only ∼1.5% of winter social connections occurring between first-order relatives [7]; therefore, relatedness was not considered a key structuring factor. Nevertheless, this social network dataset is known to be biologically meaningful, and forms part of a broader investigation into great tit social ecology, contributing insights into individual sociality [8,9], social structure and demography [7,10], and links to ecological processes such as information transmission [11,12], foraging [13, 14], breeding settlement and mating behavior [7,14,15].

All work was approved by the University of Oxford, Department of Zoology, Animal Welfare and Ethical Review Board (Approval: APA/1/5/ZOO/NASPA/Sheldon/TitBreedingEcology) and adhered to local animal research guidelines. All birds were caught, tagged, and ringed by licensed BTO ringers.

## Wytham wood mice

This data set is a trapping data set of wild rodents caught in Holly Hill, Wytham Woods 2018-2019. Rodents were trapped approximately fortnightly and new individuals tagged with a PIT-tag for permanent identification. Individuals were then released back to their point of capture. Data were collected over a 1-year period (Nov 2018–Nov 2019) from a wild population of rodents in a 4 ha (200 x 200 meters square) mixed woodland plot. Data for this study was collected as part of a longer-term rodent capture-mark-recapture study, where several rodent species were caught (*Apodemus sylvaticus*, *Apodemus flavicollis* and *Myodes glareolus*). Trapping was performed every 2-4 weeks, using 200 small folding Sherman traps (5.1 x 6.4 x 16.5cm, H. B Sherman). To ensure even trapping coverage, live traps were set with an alternating checkerboard design in every other 10 x 10 meter “Grid cell” of the 200 x 200 meters study area. Traps baited with 6 peanuts, a slice of apple and sterile cotton wool for bedding were set at dusk and collected at dawn, with all animals processed, sampled and then released inside the 100m2 grid cell they were captured in. As part of processing, captured individuals were identified to species, sexed, weighed, and aged (to juvenile or adult) based on size and pelage characteristics. At first capture, all individuals were injected subcutaneously with a passive integrated transponder tag (PIT-tag) for permanent identification. There were a few cases (∼2% of captures) where the animal died during trapping and thus was not released. This was most often due to animal being found dead in the trap (which can happen due to epidemics or other poor health especially in the spring) and in a couple of cases due to animal being put down following Schedule 1 due to extreme poor health or injury. Edges in the social network between individuals (nodes) were defined as unique mice trapped nearby (within one adjacent trap distance by Euclidean distance) in the same trapping night.

